# Transcription levels of a long noncoding RNA orchestrate opposing regulatory and cell fate outcomes in yeast

**DOI:** 10.1101/2019.12.24.887935

**Authors:** Fabien Moretto, N. Ezgi Wood, Minghao Chia, Cai Li, Nicholas M. Luscombe, Folkert J. van Werven

**Affiliations:** Cell Fate and Gene Regulation Laboratory, The Francis Crick Institute, 1 Midland Road, London, NW1 1AT, UK; Department of Cell Biology, UT Southwestern Medical Center, 6000 Harry Hines Boulevard, Dallas, TX, 75390, USA; Bioinformatics and Computational Biology Laboratory, The Francis Crick Institute, 1 Midland Road, London, NW1 1AT, UK

## Abstract

Many long noncoding RNAs (lncRNAs) act *in cis* through transcription-coupled chromatin alterations that drive changes in local gene expression. How some *cis*-acting lncRNAs promote and others repress gene expression remains poorly understood. Here we report that in *S. cerevisiae* transcription levels of the lncRNA *IRT2*, located upstream in the promoter of the inducer of meiosis gene, regulate opposing chromatin and transcription states. Low *IRT2* transcription displays enhancer RNA-like features. At these levels, *IRT2* promotes histone exchange delivering acetylated histone H3 lysine 56 to chromatin thereby facilitating recruitment of a transcription factor and consequently activating transcription. Conversely, increasing *IRT2* transcription enhances chromatin assembly and transcriptional repression. The opposing functions of *IRT2* direct a regulatory circuit, which ensures that cells expressing opposite, but not one of either, mating-type loci enter meiosis. Our data demonstrate that the transcription levels of an lncRNA are key to controlling gene expression and cell fate outcomes.

## Introduction

Long noncoding RNAs are a diverse class of RNAs typically larger than few hundred base pairs in length that have no obvious protein coding potential. These transcripts are widely expressed, constituting a large fraction of the transcriptome (Hon et al., 2017; Iyer et al., 2015; Kung et al., 2013). LncRNAs control various cellular processes including cell differentiation and stress response, and multiple roles have been implicated in diseases such as cancer (Guttman et al., 2011; Ponting et al., 2009; Wapinski and Chang, 2011). Despite extensive efforts, the function of majority of lncRNAs remains poorly understood.

Inside cells, lncRNAs play a wide range of functions by either acting *in cis* or *in trans* (Wang and Chang, 2011). A main function of *cis*-acting lncRNAs is to regulate local gene transcription. These transcripts are often generated close to protein coding genes in promoter regions or at the 3’ends of genes where they produce sense or antisense transcripts. A combination of transcription and the RNA itself subsequently modulates transcription of protein coding genes. Multiple mechanisms have been described to achieve this (Gil and Ulitsky, 2019). For example, there has been evidence that lncRNAs can form a scaffold for regulators of chromatin and transcription factors. Additionally, act of transcription is a primary mechanism of action by which some lncRNAs regulate gene expression (Kornienko et al., 2013). During lncRNA transcription, RNA polymerase II recruits a wide range of enzymes that regulate and modify chromatin, which alter the local chromatin, and thereby control gene transcription locally (Ard et al., 2017; Venkatesh and Workman, 2015). *Cis*-acting lncRNAs can have both repressive and activating functions. Examples from yeasts and other eukaryotes have demonstrated that transcription through promoters of protein coding genes exerts repression (Ard et al., 2014; Bumgarner et al., 2009; Latos et al., 2012; Martens et al., 2004; Rom et al., 2019; van Werven et al., 2012). At these loci, chromatin regulators such as facilitator of chromatin transcription (FACT), Set2-dependent deposition of lysine 36 methylation, Set3 mediated histone de-acetylation and others mediate transcriptional repression evoked by lncRNA transcription (Ard and Allshire, 2016; Hainer et al., 2011; Kim et al., 2012; van Werven et al., 2012). Conversely, lncRNA transcription can also stimulate opening of chromatin and promote coding gene transcription, which involves increased histone acetylation and disassembly of nucleosomes locally (Hirota et al., 2008; Takemata et al., 2016). In addition, production of RNAs at enhancers, also known as eRNAs, contributes to enhancer activity and thus promotes gene activation, through a mechanism that is not well understood (Li et al., 2016). Taken together, gene regulation of locally transcribed lncRNAs has a wide range of effects, from activation to repression, on coding gene expression. How transcription of some *cis*-acting lncRNAs promote and others repress gene expression remains poorly understood.

In budding yeast, intergenic lncRNA transcription is pervasive (Tisseur et al., 2011). Genes involved in metabolism, flocculation, and sporulation are directly regulated by lncRNA transcription to ensure accurate expression under the right conditions (Bumgarner et al., 2009; Hongay et al., 2006; Martens et al., 2004). One of these genes is Inducer of <u>ME</u>iosis 1, *IME1*. This master regulatory transcription factor controls the cell fate decision of whether to enter meiosis and form spores (Kassir et al., 1988; Nachman et al., 2007). Expression of *IME1* activates the so-called early meiotic genes, thereby driving meiotic entry and the production of four haploid spores (Primig et al., 2000; van Werven and Amon, 2011). The *IME1* gene is highly regulated at the level of transcription through its unusually large promoter (about 2.5 Kb), at which nutrient and mating-type signals integrate (van Werven and Amon, 2011). These signals ensure that *IME1* transcription is only induced in cells expressing opposite mating-type loci (*MAT*a and *MAT*α) under starvation conditions, in the absence of glucose and nitrogen sources. Thus, understanding how nutrient and mating-type signals integrate at the *IME1* promoter is key to understanding how yeast cells make the decision to enter meiosis.

Two lncRNAs are expressed upstream in the *IME1* promoter in tandem that mediate mating-type control of *IME1* transcription (Moretto et al., 2018; van Werven et al., 2012). In cells with a single mating type (*MAT*a or *MAT*α), typically haploid cells, the lncRNA named *IME1* regulating transcript 1 (*IRT1*) represses the *IME1* promoter to ensure that these cells do not undergo a lethal meiosis. Transcription of *IRT1* represses *IME1* by acting on a critical part of the *IME1* promoter where transcription factors important for activation of the *IME1* promoter bind (Kahana et al., 2010; Sagee et al., 1998). In *MAT*a/α diploid cells, *IRT1* transcription is reduced because the a1α2 heterodimer (expressed from opposite mating-type loci) represses the transcriptional activator of *IRT1*, *RME1*, enabling *IME1* induction and thus meiotic entry (Figure 1A) (Mitchell and Herskowitz, 1986; van Werven et al., 2012). Despite the presence of a1α2 in *MAT*a/α diploid cells, Rme1, and thus *IRT1*, is expressed to moderate levels in various genetic backgrounds (Deutschbauer and Davis, 2005; Gerke et al., 2009). To overcome *IRT1* transcription in *MAT*a/α diploid cells, Ime1 directly activates expression of a second lncRNA named *IRT2*, located upstream in its own promoter (Moretto et al., 2018). Transcription of *IRT2*, in turn, locally interferes with Rme1 recruitment, and consequently represses *IRT1* transcription and thereby promoting *IME1* expression (Figure 1A). Thus, two lncRNAs and Ime1 itself form a regulatory circuit that controls *IME1* promoter activity.

**Figure 1.**
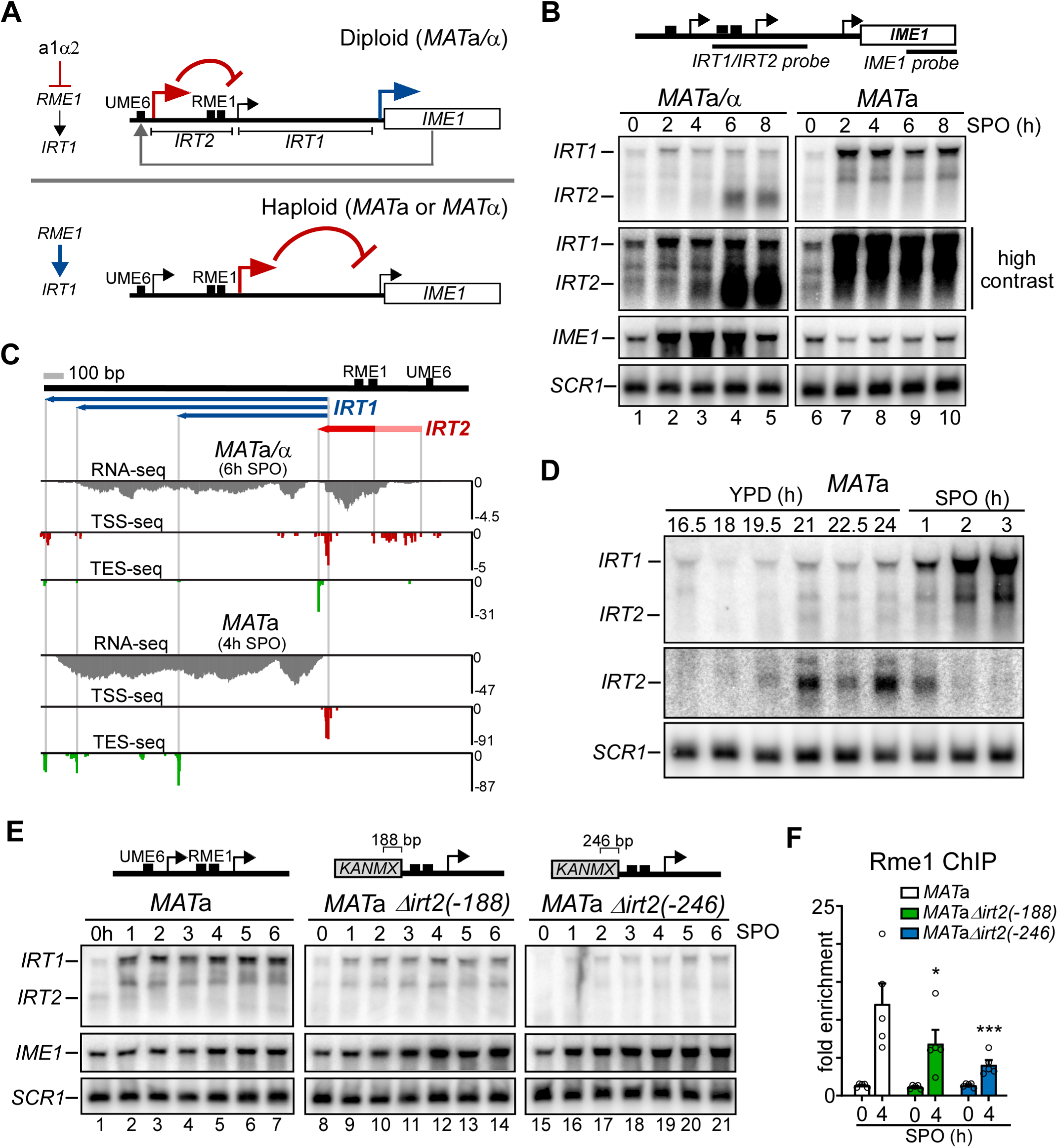
*IRT2* is required for activation of *IRT1* transcription. **A**, Schematic of the two long noncoding RNAs, *IRT1* and *IRT2*, expressed in the *IME1* promoter. In *MAT*a/α cells (diploids) expression of *RME1*, the activator of *IRT1*, is repressed by the a1α2, which allows *IME1* expression in this cell-type. A feedback regulatory circuit consisting of *IRT2*, *IRT1* and Ime1 facilitates *IME1* expression in diploids as previously described (Moretto et al., 2018). In cell with single mating-type (*MAT*a or *MAT*α, haploids), Rme1 is expressed, and *IME1* expression is repressed by transcription of *IRT1* (van Werven et al., 2012). Also depicted is the Ume6 binding site, a critical mediator for *IRT2* regulation. In the absence of Ime1, Ume6 represses *IRT2* transcription, while Ume6 interacts with Ime1 to stimulate *IRT2* transcription. **B**, *IRT1*, *IRT2* and *IME1*, expression in *MAT*a/α (lanes 1-5) (FW1509) and *MAT*a cells (lanes 6-10) (FW1511) as detected by northern blot. Cells were in grown rich medium (YPD) to saturation, shifted to pre-sporulation medium (pre-SPO), grown for an additional 16 hours, and subsequently transferred to sporulation medium (SPO). Samples were taken at the indicated time points. Northern blot membranes were probed for *IME1*, *IRT1* and *IRT2* (combined probe). *SCR1* was used as a loading control. A higher contrast for the *IRT1*/*IRT2* probed membranes is also displayed to illustrate *IRT2* signal in haploid cells. **C**, *IRT1* and *IRT2* transcription start-site sequencing (TSS-seq), transcription end site sequencing (TES-seq), RNA-seq data for *MAT*a/α (6h SPO, top panel) and *MAT*a (4h SPO, bottom panel). Cells grown as described in b. The blue lines indicate different *IRT1* RNA isoforms. Red line indicates the *IRT2* transcript, and light red line indicates where *IRT2* transcription initiates. The values on y-axes indicate Reads Per Million (RPM). **D**, Similar to b except that haploid *MAT*a cells (FW1509) were grown in rich medium to saturation for up to 24h, and directly transferred to SPO. Samples were taken at the indicated time points. To detect *IRT2* expression, an *IRT2* specific probe was also used. **E**, *IRT1*, *IRT2*, and *IME1* expression in WT *MAT*a cells (FW1509, lanes 1-7), and Δ*irt2*(−188) (FW1210, lanes 8-14) and Δ*irt2*(−246) (FW128, lanes 15-21) mutants. Gene deletions were generated using a one-step deletion protocol resulting in 188 and 246 base pairs (bp) of *IRT2* deleted, while keeping the Rme1 binding sites in *IRT2* intact. Cells were grown as described in b. *SCR1* was used as loading control. Samples were taken at the indicated time points. **F**, Rme1 association to the *IRT1* promoter in mutants described in d. These cells also harboured *RME1* tagged with V5 epitope (FW4031, FW3132, and FW3140). Cells were grown and treated as described in b, samples were taken at the indicated time point, and fixed with formaldehyde. Chromatin immunoprecipitation was performed, and recovered DNA fragments were quantified by quantitative PCR using a primer pair directed to Rme1 binding sites in the *IRT1* promoter. Signals were normalized to *HMR*, which does not bind Rme1. The error bars represent the standard error of the mean (SEM) of n = 5 biological repeats. * and *** correspond to a p-value < 0.05 and < 0.005 compared to *MAT*a control on a two-way ANOVA followed by a Fisher’s LSD test.

The nutrient and mating-type signals act in hierarchical order to control the decision to enter meiosis. In a rich nutrient environment, the *IME1* promoter is repressed independently of the mating-type status of the cell. However, cells exposed to starvation signals display mating-type dependent expression of *IME1*. Under starvation conditions, *MAT*a/α diploid cells induce *IME1* transcription and subsequently enter meiosis, while cells with a single mating-type induce *IRT1* transcription to ensure that the *IME1* promoter remains repressed (Moretto and van Werven, 2017). Thus, the mating-type status of the cell determines the outcome of *IME1* and *IRT1* transcription states under starvation conditions.

Here we describe the mechanism by which yeast cells ensure that mating-type control of the *IME1* promoter is activated in a timely and robust manner. We demonstrate a dual role for the more upstream lncRNA in the *IME1* promoter, *IRT2*. In short, we find that transcription levels of *IRT2* regulate opposing chromatin and transcription states. In cells with a single mating-type low levels of *IRT2* promotes expression of the adjacent lncRNA *IRT1* by directing H3 lysine 56 acetylation to chromatin. By contrast, increasing *IRT2* transcription drives chromatin assembly and repression of *IRT1*. The dual function of *IRT2* shapes the regulatory circuit, which ensures that cells expressing opposite, but not one of either, mating-type loci (*MAT*a and *MAT*α) enter meiosis. Taken together, our data demonstrates that transcription of an lncRNA, even at low levels, can play a critical role in regulating gene expression, and that changes in lncRNA transcription levels can lead to distinct regulatory and cell fate outcomes.

## Results

### *IRT2* is required for repression of *IME1* in cells with a single mating-type

We hypothesized that there is a feedback signal that ensures a robust transition from nutrient to mating-type control of *IME1* promoter activity, possibly involving Ime1 itself and *IRT2*. To examine whether there is such a function for *IRT2*, we first measured the expression of *IRT1* and *IRT2* and mapped the transcription start and polyadenylation sites (TSS and TES) of both transcripts in cells harbouring single mating-type (*MAT*a) and both mating-types (*MAT*a/α). As expected, *MAT*a/α diploid cells entering meiosis synchronously, induced *IME1* expression rapidly, displayed strong induction of *IRT2*, while *IRT1* levels remained relatively low (Figure 1A, 1B and S1A) (van Werven et al., 2012). In these cells (6h SPO, Figure 1C and S1B), we detected a single *IRT2* polyadenylation site (TES-seq) while multiple transcription start sites (TSS-seq) spread over ∼ 215 base pairs (bp) were matching the slight smear observed on the northern blot for the *IRT2* transcript (Figure 1B). In haploid cells (*MAT*a), *IRT1* expression was much higher than in *MAT*a/α cells and *IME1* expression was repressed (Figure 1A and 1B). The *IRT1* transcription start site mapped to a single region while one polyadenylation site (TES-seq) mapped to the middle of *IRT1* and two sites were detected near the 3’end (Figure 1C and Figure S1B). In line with this observation, two distinct *IRT1* species were detected by northern blot, which were reduced to one truncated transcript when *IRT1* transcription was terminated early (*irt1-T)* (Figure 1B (lanes 7-10) and Figure S1C). Surprisingly, we also detected low levels of *IRT2* expression prior to *IRT1* induction in *MAT*a cells (Figure 1B (see lane: 6)). The *IRT2* transcript (RNA-seq), start and end sites (TSS-seq and TES-seq), were also detectable at low levels in starved *MAT*a cells (SPO 4h) demonstrating that *IRT2* is also expressed in this cells type (Figure S1D). In order to capture the *IRT2* expression window in haploid *MAT*a cells, we sampled over a prolonged period of time before induction of starvation (Figure S1A). Strikingly, we detected *IRT2* expression in several time points prior to *IRT1* induction (Figure 1D). To test whether *IRT2* is required for *IRT1* induction, we created partial deletions disrupting the *IRT2* TSS cluster (Δ*irt2*(−188) and Δ*irt2*(−246)) while keeping the Rme1 binding sites intact (Figure 1E and S1E). Remarkably, in Δ*irt2*(−188) and Δ*irt2*(−246) cells *IRT1* expression decreased, *IME1* levels increased, and as expected *IRT2* expression was not detectable. Both *IRT2* truncation mutants also displayed reduced association of Rme1 to the *IRT1* promoter (Figure 1F). Thus, in addition to the transcriptional repressor function described for *IRT2* in *MAT*a/α cells previously (Moretto et al., 2018), *IRT2* is also required for *IRT1* expression and thus repression of the *IME1* gene in cells with single mating-type (*MAT*a or *MAT*α) cells.

### Transcription of *IRT2* prevents meiotic entry in cells with a single mating-type

We next evaluated whether transcription of *IRT2* contributes to *IRT1* activation. First, we integrated a transcriptional terminator between the *IRT2* transcription start site and the Rme1 binding sites to generate the *irt2-T* allele (Figure 2A). A shorter form of *IRT2* was detected in *irt2-T* cells (*IRT2*\*, Figure 2A). Remarkably, *irt2-T* cells showed diminished association of Rme1 with the *IRT1* promoter, reduced *IRT1* expression and RNA polymerase II (Pol II) binding to *IRT1*, and increased *IME1* expression (Figure 2A, 2B and 2C). A control sequence of about the same size as the transcriptional terminator sequence (*irt-I*) did not alter *IRT1* and *IME1* expression (Figure S2A and S2B (see lanes: 2-5 and 12-15)). In addition, a transcriptional terminator integrated into *IRT1* (*irt1-T*) showed wild-type like Rme1 recruitment and displayed no additional reduction in Rme1 binding when combined with Δ*irt2*(−246), indicating that transcription through *IRT1* is not required for inducing *IRT1* expression (Figure S2C). Second, we examined whether RNA polymerase II (Pol II) plays role in activation of *IRT1* by transiently depleting a core subunit, Rpb3, prior to *IRT1* induction using an auxin inducible degron (Rpb3-AID) (Figure 2F and Figure S2D (see lanes: 2-3)) (Nishimura et al., 2009). While transient Rpb3-AID depletion did not affect Rme1 protein expression, Rme1 association with the *IRT1* promoter was reduced suggesting that transcription by Pol II is required prior to Rme1 recruitment (Figure 2D, S2D, and S2E). Third, we modulated *IRT2* transcription without affecting the *IRT2* sequence. We reasoned that if *IRT2* transcription was involved in *IRT1* activation then changing *IRT2* expression should affect the *IRT1* expression pattern. Therefore, we first abrogated *IRT2* activation by introducing point mutations in Ime1 (*ime1-T*), which impairs Ime1 function (Moretto and van Werven, 2017). Little or no *IRT1* expression was detected in the absence of *IRT2* transcription (Figure 2E and 2F). Additionally, we constitutively expressed *IRT2* by deleting the Ume6 repressor binding site (*u6bs*Δ) in the *IRT2* promoter, which decouples *IRT2* expression from *IME1* activation (Figure 2E) (Moretto et al., 2018). To confirm that *u6bs*Δ gives rise to *IRT2* transcription, we mapped the transcript (Figure 2G). As expected, the *IRT2* start and end sites in *u6bs*Δ overlapped with positions we identified for *IRT2* (Figure 1C and Figure S1D). Constitutive levels of *IRT2* transcription (*u6bs*Δ) led to earlier *IRT1* transcription and earlier Rme1 recruitment (Figure 2H (see lanes: 1-5 and 9-13), Figure S2F and S2G (see lanes: 1-8, 12-19, and 23-30) and S2H). Furthermore, *u6bs*Δ rescued the *IRT1* expression defect observed when Ime1 function was impaired (*ime1-T*), but not when *IRT2* transcription was terminated earlier (*irt2-T)* (Figure S2I (see lanes: 7-9 and 10-12) and S2J (see lanes: 12-15 and 17-20)). We conclude that Ime1 activates *IRT1* expression via *IRT2* transcription. Taken together, these data demonstrate that transcription of *IRT2* is required for induction of *IRT1* expression in cells with a single mating-type.

**Figure 2.**
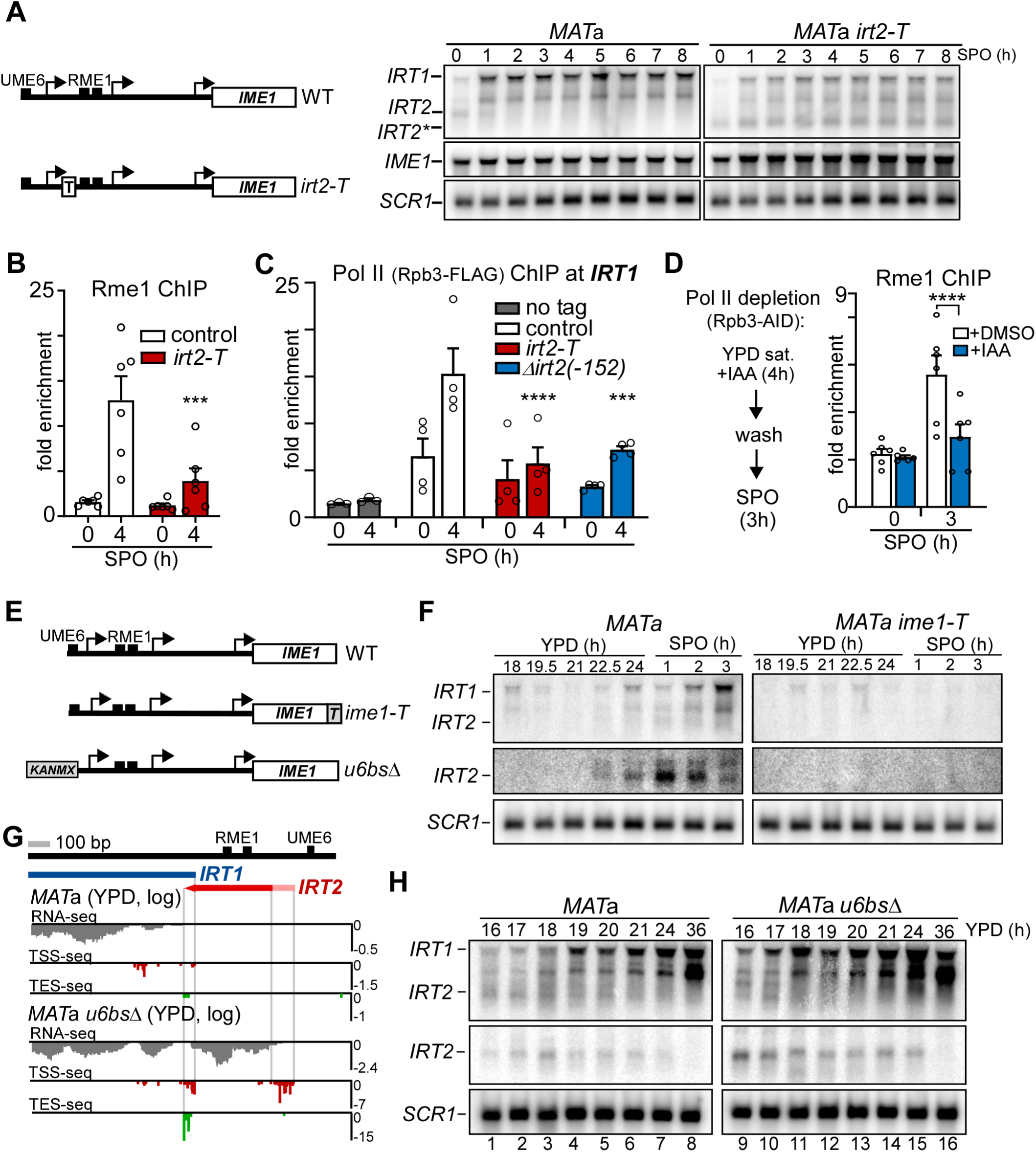
Transcription of *IRT2* is required for induction of *IRT1* expression. **A**, A schematic of *IME1* promoter harbouring a transcriptional terminator which was integrated between the *IRT2* transcription start site and the Rme1 binding sites, *irt2-T* (left); *IRT1*, *IRT2*, and *IME1* expression in WT and *irt2-T MAT*a cells (FW1509 and FW3596) as detected by northern blot. Cells were grown in rich medium to saturation, shifted and grown in pre-SPO, and subsequently transferred to SPO. The asterisk depicts a short form of *IRT2* caused by early termination in *IRT2*. *SCR1* was used as a loading control (right). **B**, Rme1 association at the *IRT1* promoter in WT and *irt2-T MAT*a cells (FW4031 and FW3128) by ChIP of Rme1-V5. Recovered DNA fragments were quantified by qPCR using a primer pair directed to Rme1 binding sites. Signals were normalized to *HMR*, which does not bind Rme1. The error bars represent the SEM of n = 6 biological repeats. *** correspond to a p-value < 0.005 respectively on a two-way ANOVA followed by a Fisher’s LSD test. **C**, RNA polymerase II (Pol II) association at the *IRT1* in *MAT*a (control), *irt2-T* and Δ*irt2*(−246) (FW8515, FW8512 and FW8510 respectively) in cells expressing FLAG-tagged Rpb3 at 0h and 4h in SPO. A no tag control (FW1509) was included for the analyses. Error bars represent the SEM of n = 4 biological repeats except for the no tag condition n = 3. *** and **** correspond to a p-value < 0.0005 and < 0.0001 respectively on a two-way ANOVA followed by a Fisher’s LSD test performed on the whole group of samples including the one presented in Figure S2E. **D**, Schematic for depleting RNA polymerase using Rpb3 auxin induced degron (AID). Cells were grown to saturation in rich medium (YPD) treated with IAA to deplete Pol II, subsequently IAA was washed out from the medium, and cells were transferred to SPO (left). ChIP of Rme1 using *MAT*a cells harbouring *RPB3-AID* and *RME1-V5* (FW6467). **** correspond to a p. value < 0.0001 on a two-way ANOVA followed by Fisher’s LSD test. **E**, Schematic of *ime1-T* and *u6bs*Δ mutants. The *ime1-T* mutant harbours point mutations in the C-terminus of *IME1*, which impairs Ime1 function. The *u6bs*Δ harbours a deletion of the Ume6 binding site, which is located in the *IRT2* promoter and is critical for regulating *IRT2* transcription. **F**, *IRT1* and *IRT2* expression in WT and *ime1-T MATa* cells (FW1509 and FW2189) detected by northern blot. Cells were grown in rich medium (YPD) to saturation, shifted to SPO, and samples were taken at the indicated time points. *SCR1* was used as a loading control. **G**, *IRT2* transcript start and end sites in *u6bs*Δ cells determined by TSS-seq and TES-seq during exponential growth in YPD. The values on y-axes indicate Reads Per Million (RPM). **H**, Same as F, except that WT (FW1509, lanes 1-8) and *u6bs*Δ *MATa* cells (FW2438, lanes 9-16) were used and cells were not shifted to SPO.

Mis-expression of meiotic genes can have detrimental consequences to haploid cells (Lino and Yamamoto, 1985; Wagstaff et al., 1982). Haploid cells harbouring a single mating-type, but lacking Rme1 and *IRT1*, undergo a lethal type of meiosis (van Werven et al., 2012). Therefore, we determined the importance of *IRT2* mediated activation of *IRT1* in preventing haploid cells from entering meiosis. We found that in *irt2* mutants (Δ*irt2*(−246) and *irt2-T*), a large fraction of cells displayed high levels of *IME1* expression (more than 30 mRNA copies per cell) (Figure 3A, S3A, and S3B). After a prolonged period of starvation, *irt2* mutants (Δ*irt2*(−188), Δ*irt2*(−246) and *irt2-T*) also displayed reduced viability possibly because these cells underwent lethal meiosis (Figure 3B). To examine the effects on meiosis further, we generated diploid cells of these *irt2* mutants harbouring a single mating type (*MAT*a/a) thus mimicking mating-type repression of *IME1* expression. Approximately 30 to 40 percent of *MAT*a/a diploid cells for each *irt2* mutant underwent at least one meiotic division (Figure 3C). We conclude that *IRT2* is essential for inhibiting meiotic entry in starved cells with a single mating-type.

**Figure 3.**
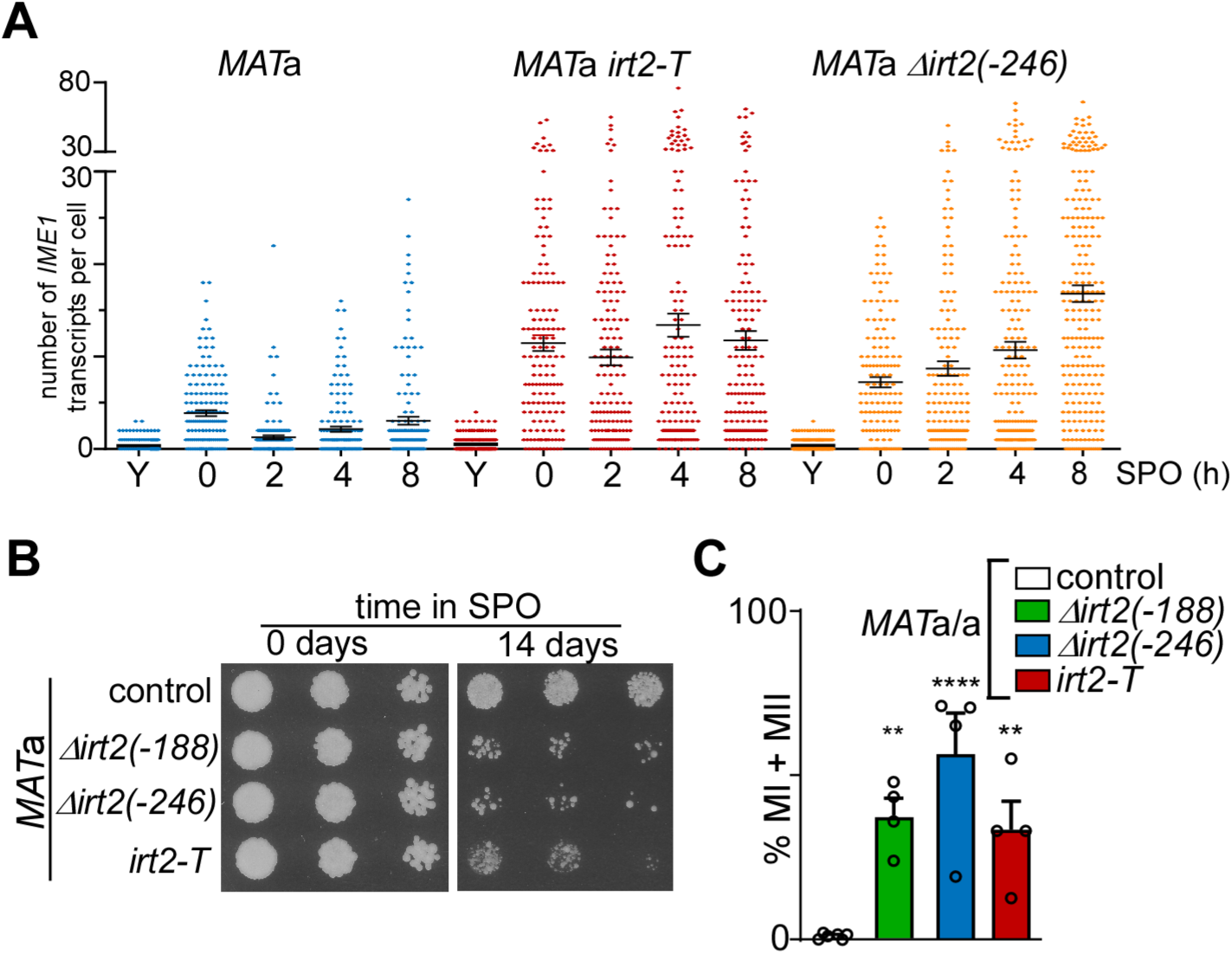
Transcription of *IRT2* prevents meiotic entry in cells with a single mating-type. **A**, *IME1* expression in single cells as measured by single-molecule RNA FISH. Cells were grown for 24h to saturation in rich medium (Y), transferred and grown in pre-SPO, and shifted to SPO. Cells were fixed at the indicated time points and hybridized with probes directed against *IME1* and *ACT1*. Each dot represents the number of *IME1* transcripts in a single cell positive for *ACT1* expression. The mean and SEM are also depicted. n=150 cells. **B**, Spot assay after cells were exposed for 0 or 14 days in SPO. In short, WT, Δ*irt2(−188)*, Δ*irt2(−246)*, and *irt2-T* cells (FW1509, FW1210, FW128 and FW3596) were grown in rich medium, shifted to pre-sporulation medium, and subsequently transferred to SPO for 0 and 14 days before cells were spotted on rich medium agar plates in five-fold, serial dilutions. **C**, *IRT2* prevents entry into meiosis in *MAT*a/a diploid cells. *MAT*a/a diploid control cells (FW15) and cells harbouring Δ*irt2(−188)*, Δ*irt2(−246)* or *irt2-T* (FW3453, FW3402, and FW3629) were grown as described in I. After 72 hours in SPO cells were fixed, and DAPI masses were counted. Cells harbouring 2, 3 or 4 DAPI masses were considered to have undergone meiotic divisions (MI+MII). n = 4 +/-SEM except for the control sample n = 6. ** and **** correspond to a p-value < 0.005 and < 0.0001 respectively, on a one-way ANOVA followed by Fisher’s LSD test.

### *IRT2* transcription directs H3K56ac to chromatin to promote *IRT1* activation

Transcription of *IRT2* displays features of enhancer RNAs (eRNAs) in mammalian cells (Li et al., 2016). Production of eRNAs stimulates recruitment of transcription factors and promotes enhancer activity. With this view, *IRT2* stimulates transcription factor recruitment and activation of the downstream lncRNA *IRT1*. Transcription of eRNAs also controls histone modifications including histone acetylation, a mark of active transcription (Bose et al., 2017). To gain insight into the mechanism, by which *IRT2* stimulate *IRT1* expression, we screened for mutants that displayed decreased *IRT1* and increased *IME1* expression. We specifically focussed on a small set of known histone modifying enzymes as well as factors that are part of the Pol II machinery. Gene deletions affecting Pol II transcription fidelity (*RPB9* and *CTK1*), histone acetylation (*GCN5* and *RTT109*), and histone chaperone function (*ASF1*) were identified (Figure 4A and Table S1). We focussed our analyses on two candidate genes: *RTT109* and *ASF1*. Rtt109 is the sole histone acetyltransferase that acetylates histone H3 lysine 56 (H3K56ac) in yeast, whereas Asf1 is involved directly in chromatin assembly and acts as a chaperone for Rtt109 directed H3K56ac (Driscoll et al., 2007; Masumoto et al., 2005; Recht et al., 2006; Schneider et al., 2006; Tsubota et al., 2007). H3K56ac marked histones are assembled into nucleosomes during DNA replication where it buffers gene dosage imbalances, but is also present at promoters where it is incorporated into nucleosomes in a replication-independent manner (Kaplan et al., 2008; Rufiange et al., 2007; Schneider et al., 2006; Voichek et al., 2016; Williams et al., 2008; Xu et al., 2005). Furthermore, nucleosomes harbouring H3K56ac mark active transcription and active enhancers in higher eukaryotes (Schneider et al., 2006; Skalska et al., 2015; Varv et al., 2010). We found that in *rtt109*Δ, and to a lesser extent in *asf1*Δ *MAT*a cells, *IRT1* expression was reduced and *IME1* expression levels increased (Figure 4B and S4A). Importantly, Rme1 protein levels were not affected in *rtt109*Δ cells (Figure S4B). In addition to steady-state RNA measurements, we also determined whether *IRT1* and *IME1* transcription (Pol II ChIP and nascent RNA-seq) was affected in *rtt109*Δ cells (Figure 1C, 1D, S4C and S4D). *rtt109*Δ cells displayed reduced *IRT1* transcription, less Pol II binding to *IRT1* and increased *IME1* transcription. Further genetic analysis suggested that Rtt109 acts downstream of *IRT2* transcription. First, *rtt109*Δ combined with early termination of *IRT2 (irt2-T)* displayed a defect in *IRT1* expression and Rme1 association with the *IRT1* promoter that was comparable to that of the single mutants (Figure 4E (see lanes: 6-10, 11-15 and 16-20), 4F and Figure S4E). Second, de-repression of *IRT2* transcription (*u6bs*Δ) did not rescue the *IRT1* expression defect in *rtt109*Δ cells (Figure S4F (see lanes: 6-10, 11-15 and 16-20), and S4G). It is also worth noting that the *IRT2* RNA was clearly detectable in *rtt109*Δ *u6bs*Δ cells suggesting that the *IRT2* RNA by itself has little function in *IRT2*-mediated activation of *IRT1* (Figure S4F (see lane: 16), and S4G).

**Figure 4.**
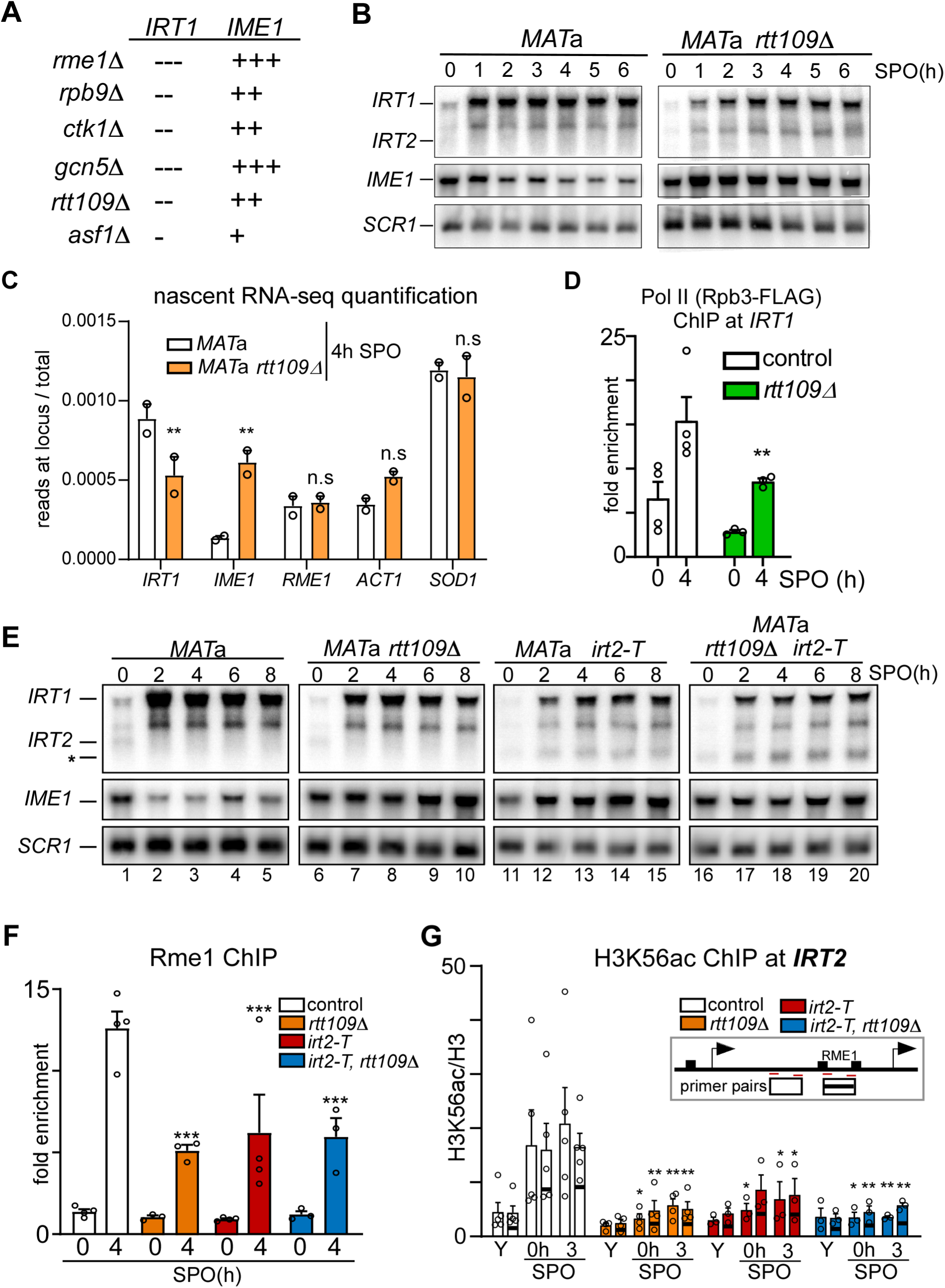
Rtt109 and histone H3 lysine 56 acetylation mediate *IRT2* directed activation of *IRT1* transcription. **A**, Summary of candidate genes involved in *IRT2* mediated activation of *IRT1*. The minus symbol “-“ means lower *IRT1* expression compared to *MAT*a WT cells, while the plus symbol “+” means higher *IME1* expression compared to *MAT*a WT cells. **B**, *IRT1*, *IRT2*, and *IME1* expression in WT *MAT*a cells (FW1509) and *rtt109*Δ (FW4077) cells as detected by northern blot. *SCR1* was used as loading control. Cells were grown in rich medium to saturation, shifted and grown in pre-SPO, and subsequently transferred to SPO. Samples were taken at the indicated time points. **C**, Quantification of the reads associated with the selected transcripts (*IRT1*, *IME1* and the controls *RME1*, *ACT1* and *SOD1*) as obtained during nascent RNA-seq described in f. Raw reads associated with the different loci were sum up and a ratio over the total aligned reads for the sample was calculated. n = 2 +/-SEM. ** correspond to a p-value < 0.005 on a parametric unpaired two-tailed Student’s *t*-test. **D**, Pol II association with *IRT1* in control *MAT*a and *rtt109*Δ cells (FW8515, FW8561). These cells also harbored Rpb3-FLAG, which was used for the immunoprecipitation step. Cells were grown as described in b, and cells were formaldehyde fixed at the indicated time points. The silent mating type locus *HMR* was used to normalize the signals. SEM and n >= 3. ** correspond to a p-value < 0.005 on a two-way ANOVA followed by a Fisher’s LSD test performed on the whole group of samples including those presented in figure 2d. **E**, Expression of *IRT1*, *IRT2*, and *IME1* in *MAT*a WT cells (FW1509, lanes 1-5), compared to *rtt109*Δ (FW4077, lanes 6-10), *irt2-T* (FW3596, lanes 11-15), single and *rtt109*Δ, *irt2-T* double mutants (FW4072, lanes 16-20). Cells were grown in rich medium (YPD) to saturation, shifted and grown in pre-SPO, and subsequently transferred to SPO. Sample were taken at the indicated time points. *SCR1* was used as a loading control. **F**, ChIP of Rme1-V5 at the *IRT1* promoter in mutants described in e, but harbouring the *RME1-V5* allele (FW4031, FW3128, FW4075, and FW4073). n = 4 +/-SEM for control and *irt2-T* and n = 3 for *rtt109*Δ and *irt2-T rtt109*Δ. *** correspond to a p-value < 0.0005 on a two-way ANOVA followed by Fisher’s LSD test. **G**, Histone H3 lysine 56 acetylation (H3K56ac) levels in the *IRT1* promoter as measured by ChIP in control (*MAT*a, FW1509) and *irt2-T* (FW3596) cells. Samples were taken during exponential growth (YPD), pre-SPO (0h), and SPO (3h). H3K56ac ChIP signals were normalized to histone H3. As control *rtt109*Δ and *irt2-T rtt109*Δ cells (FW4077, and FW4072) were included in the analyses. n = 5 +/-SEM for control, n = 4 for *rtt109*Δ and n = 3 for *irt2-T* and *irt2-T rtt109*Δ. * and ** correspond to a p-value < 0.05 and < 0.005 respectively on a two-way ANOVA followed by Fisher’s LSD test performed on each primer pair individually.

Next, we evaluated whether H3K56ac is responsible for *IRT2* mediated activation of *IRT1*. In actively transcribed gene bodies, histone H3 lysine 36 methylation (H3K36me) is known to limit the incorporation of H3K56ac into nucleosomes (Venkatesh et al., 2012). In previous work, however, we showed that H3K36me is absent in the *IRT2* region of the *IME1* promoter even during active *IRT2* transcription (Moretto et al., 2018). Therefore, transcription of *IRT2* might allow H3K56ac incorporation into nucleosomes. Indeed, when we measured H3K56ac levels in the *IRT2* region, we found that H3K56ac was enriched at the time of *IRT2* transcription (0h and 3h), but not when *IRT2* was repressed during exponential growth (YPD), or when *RTT109* was deleted (Figure 4G). Importantly, H3K56ac levels were reduced in *irt2-T* cells further supporting that *IRT2* transcription is required for H3K56ac deposition (Figure 4G). This data suggest that H3K56ac is directly involved in IRT2 mediated activation.

Although the main substrate of H3K56, Rtt109 has also been shown to acetylate lysine 9 of histone H3. Therefore, to examine the role of H3K56ac directly, we mutated the H3K56 residue to alanine or arginine (H3K56A and H3K56R) to mimic the absence of H3K56ac in cells. The H3K56R mutant, and to a lesser extent H3K56A, displayed reduced *IRT1* expression, and increased *IME1* expression (Figure 5A (see lanes: 1-5 and 11-15) and Figure S5A). Importantly, *IRT1* expression levels in the H3K56R mutant were affected to a degree comparable to that of the *rtt109*Δ H3K56R double mutant, suggesting that other targets of Rtt109 do not play a major role in *IRT2* mediated activation of *IRT1* (Figure S5B (see lanes: 14-16, 10-12 and 6-8) and S5C). We also examined whether H3K56ac was necessary to suppress meiotic entry in cells with single mating-type. Indeed, approximately 20 to 30 percent of *MAT*a/a diploid cells underwent at least one meiotic division when H3K56ac deposition was impaired (Figure 5B). These data demonstrate that Rtt109-mediated deposition of H3K56ac is critical for *IRT2* mediated activation of *IRT1* and prevention of inappropriate meiotic entry.

**Figure 5.**
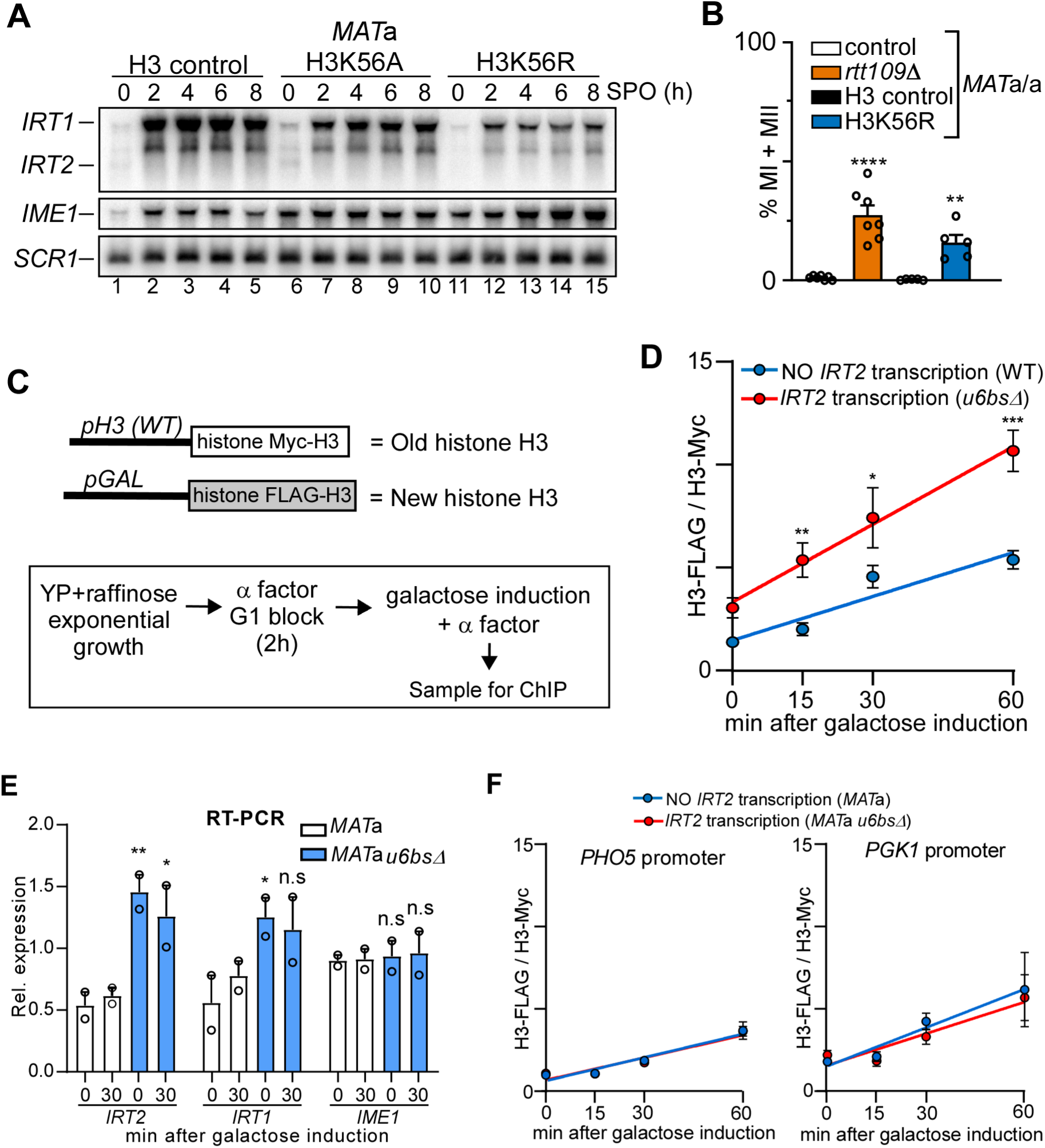
*IRT2* links H3K56ac to nucleosome dynamics. **A**, Expression of *IRT1*, *IRT2* and *IME1* in the *MAT*a histone H3 control (FW5102, lanes 1-5), H3K56A (FW5113, lanes 6-10) and H3K36R (FW5116, 11-15) cells. Cells were grown in rich medium (YPD) to saturation, shifted and grown in pre-SPO, and subsequently transferred to SPO. Sample were taken at the indicated time points. *SCR1* was used as a loading control. **B**, H3K56ac is required for preventing entry into meiosis in *MAT*a/a diploid cells. *MAT*a/a diploid cells harbouring *rtt109*Δ, H3K36R and matching controls (FW15, FW4557, FW7413 and FW7417). Cells were grown as in a. After 72 hours in SPO cells were fixed, and DAPI masses were counted. Cells harbouring 2, 3 or 4 DAPI masses were considered meiotic (MI+MII). n = 7 +/-SEM for control and *rtt109*Δ and n = 5 for H3 control and H3K56R. ** and **** correspond to a p-value < 0.005 and < 0.0001 respectively, on an unpaired parametric two-tailed Student *t*-test comparing mutant with respective control. **C**, Scheme of strain used to measure histone H3 exchange rates and experimental set up. **D**, Histone H3 exchange rate at the *IRT1* promoter in the presence or absence of *IRT2* transcription. A strain harbouring differentially epitope-tagged histone H3, with one copy expressed from the endogenous promoter (Myc-H3) and the other expressed from a *GAL1-10* inducible promoter (*pGAL-FLAG-H3*) was used for the analysis. To have constitute low levels of *IRT2* transcription we used the *u6bs*Δ (FW7880), while wild-type cells (FW7853) display no *IRT2* transcription in rich medium (YP). Cells were grown till mid-log in YP plus raffinose, arrested in G1 with **α** factor, and FLAG-H3 was induced with galactose. Samples were taken at the indicated time point for ChIP. The signals for each histone H3 ChIP (Myc-H3 and FLAG-H3) were normalized to a telomere locus, and ratio for n = 3 +/-SEM is displayed. *, ** and *** correspond to p-value < 0.05, < 0.005 and < 0.0005 respectively, on a two-way ANOVA followed by Fisher’s LSD test. The slopes of the linear regression equations (Y = 0.07105*X + 1.466 for control and Y = 0.1258*X + 3.325 for *u6bs*Δ) are significantly different. **E**, Relative expression of *IRT2*, *IRT1* and *IME1* in cells described in D (FW7880 and FW7853). Cells were grown till mid-log in YP plus raffinose, arrested in G1 with α factor, and H3-FLAG was induced with galactose. Samples were taken at the indicated time points. RNA was subjected to reverse transcription and quantitative PCR. Signals were normalized to *ACT1*. n = 2 +/-SEM. * and ** correspond to p-value < 0.05 and < 0.005 respectively on an unpaired Student’s *t*-test. **F**, Similar as D, Histone exchange at the *PHO5* and *PGK1* promoters. Strains as described in D (FW7880 and FW7853). Cells were grown as described in D. Samples were taken at the indicated time point for ChIP. Primer pairs for the *PHO5* and *PGK1* promoter were used for the analysis. The signal for each histone H3 ChIP (Myc-H3 and FLAG-H3) were normalized to a telomere locus, and ratio for n = 3 +/-SEM is displayed. The slopes of the linear regression equations for both loci are not significantly different.

How is H3K56ac incorporated into chromatin by *IRT2* transcription? The act of transcription can deliver free histones to nucleosomes in exchange for old ones (Das and Tyler, 2013; Jackson, 1990; Venkatesh and Workman, 2015). As Rtt109 mediated H3K56 acetylation occurs off chromatin, we speculated that transcription mediated histone exchange delivers H3K56ac to chromatin (Tsubota et al., 2007). To examine whether *IRT2* promotes incorporation of new histones, we measured histone H3 exchange in the presence or absence of *IRT2* transcription in a replication independent manner. A strain harbouring differentially epitope tagged histone H3, with one copy expressed from the endogenous promoter and the other expressed from a *GAL1* inducible promoter was used for the analysis (Figure 5C) (Schermer et al., 2005). Remarkably, the rate of incorporation of newly synthesized histone H3 vastly increased in the presence of constitutive levels of *IRT2* transcription (Figure 5D, compare WT (no *IRT2* transcription) to *u6bs*Δ (*IRT2* transcription)). Galactose induction had no effect on *IME1* expression under these conditions, which excludes the possibility that the effects were due to changes in *IME1* promoter activity (Figure 5E). As expected, the histone H3 exchange rate at two control promoters was not affected by *IRT2* transcription (Figure 5F and Figure S5D). We propose that in haploid cells, transcription of the lncRNA *IRT2* stimulates histone exchange, which directs H3K56ac to chromatin. Subsequently, the presence of the H3K56ac mark facilitates chromatin disassembly, recruitment of Rme1, and activation of *IRT1* transcription.

### Different transcription levels of *IRT2* regulate opposing chromatin states

Our data demonstrates that low levels of *IRT2* transcription activates *IRT1* expression in haploid cells harbouring a single mating-type (*MAT*a or *MAT*α). Previously, we showed that transcription of *IRT2* is important for repressing *IRT1* in *MAT*a/α diploid cells, which promotes *IME1* expression and meiotic entry (Moretto et al., 2018). In order to reconcile the two seemingly contradictory observations, we controlled *IRT2* transcription to high levels by replacing the endogenous promoter with the *CUP1* promoter (*pCUP-IRT2*) in *MAT*a/a diploids (which behave like *MAT*a haploid cells) (Figure 6A). *MAT*a/a cells did not undergo meiotic divisions because *IRT2* expression from the *CUP1* promoter is relative low under noninducing conditions (-Cu) and thus promotes *IRT1* transcription as in wild-type cells (Moretto et al., 2018). When we induced *IRT2* transcription to high levels (+Cu), a large fraction of cells underwent meiosis, which strongly suggests that *IRT1* transcription was repressed. These data are consistent with our findings reported previously (Moretto et al., 2018), and indicate that different *IRT2* transcription levels have opposite effects on *IRT1* expression.

**Figure 6.**
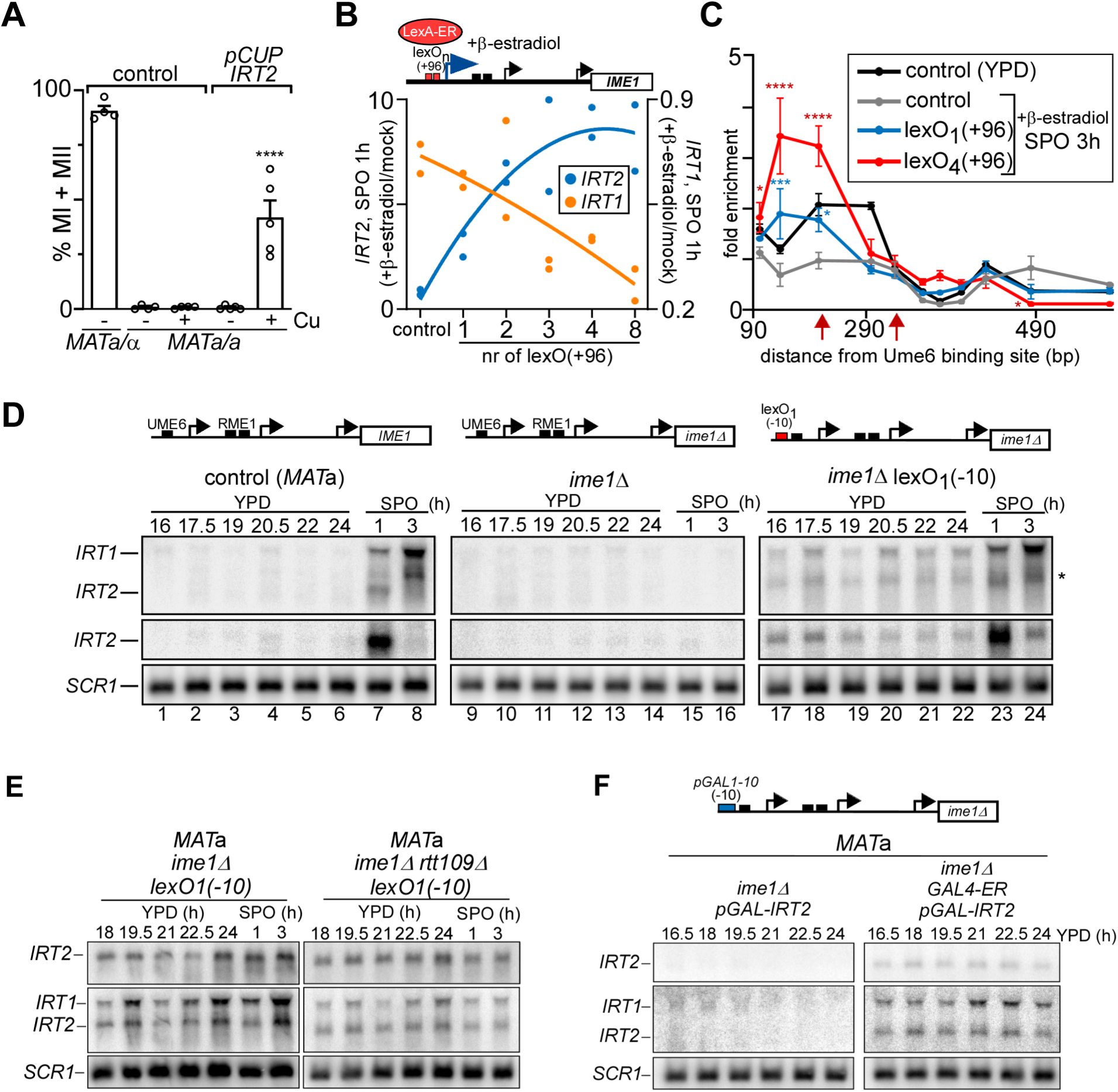
Distinct levels of *IRT2* lead to opposite outcomes of *IRT1* transcription. **A**, High levels of *IRT2* transcription promotes entry into meiosis in *MAT*a/a diploid cells. *MAT*a/α and *MAT*a/a diploid cells (FW1511 and FW15) and *MAT*a/a cells harbouring *pCUP-IRT2* (FW8923) were shifted to SPO and either treated (+Cu) or not (-Cu) with copper sulphate. After 72 hours in SPO cells were fixed, and DAPI masses were counted. Cells harbouring 2, 3 or 4 DAPI masses were considered to have undergone meiotic divisions (MI+MII). n = 4 +/-SEM for the controls and n = 5 for *pCUP-IRT2*. **** correspond to a p-value < 0.0001 on a two-way ANOVA followed by Fisher’s LSD test performed on *MAT*a/a strains without or with copper sulphate treatment. **B**, Increasing *IRT2* transcription levels leads to increasing repression of *IRT1* expression. Scheme for controlling different levels of *IRT2* by LexA-ER (top). Multiple lexA operator (lexO) sequences were integrated in the *IRT2* promoter at +96 bp relative to *IRT2* start site (lexO(+96)). LexA-ER is activated by β-estradiol. *IRT1* and *IRT2* levels as detected by northern blot and normalized to *SCR1* (bottom), in cell harbouring 1, 2, 3, 4, and 8 lexO(+96) sites in the *IRT2* promoter (FW6594, FW6599, FW6607, FW6611 and FW6619) at 1 hours in SPO. The *MAT*a LexA-ER control strain (C, FW6560) was included in the analysis. Cells were grown in rich medium till saturation, shifted to pre-SPO medium, treated (+β-estradiol) or not treated (mock) for 3 h, and shifted to SPO plus β-estradiol. The ratio of +β-estradiol/mock is displayed. n=2 data points and a trend line representing second degree polynomial fit are shown. **C**, Chromatin structure at the *IRT1* promoter in the presence of distinct levels of *IRT2* transcription. Control cells (*MAT*a LexA-ER, FW6560) or cells harbouring 1 or 4 lexO(+96) sites (FW6594 or FW6611) were grown as described in b, cell were fixed with formaldehyde, and chromatin were digested with micrococcal nuclease (MNase) followed by qPCR using scanning primer pairs in *IRT2*. The red arrows indicate the position of the Rme1 binding sites. The signals were normalized over a telomere locus. n = 3 +/-SEM. *, *** and **** correspond to a p-value < 0.05, < 0.0005 and < 0.0001 respectively, on a two-way ANOVA followed by Fisher’s LSD test performed on lexO strains compared to control SPO 3h. **D**, *IRT2* expression from a single lexO site integrated at -10 bp (lexO_1_(−10)) in the *IRT2* promoter rescues the *IRT1* expression defects of the *ime1*Δ strain. *MAT*a WT cells (FW1510, lanes 1-8), or cells harbouring *ime1*Δ together with the WT *IRT2* promoter (FW1556, lanes 9-16) or a lexO_1_(−10) site (FW7142, lanes 17-24) were grown in YPD to saturation, and shifted to SPO. Samples were taken at the indicated time points. *SCR1* was used as a loading control. The asterisks labelled band represents a longer version of *IRT2* expression from lexO_1_(−10) and truncated version *IRT1*, which have about same size. **E**, *IRT1* and *IRT2* expression in *MAT*a *ime1*Δ cells harboring one copy of lexO integrated -10 bp from *IRT2* start site (lexO_1_(−10)) (FW7142), or the same cells also containing *rtt109*Δ (FW8555). Cells were grown as described in D. Samples were taken at the indicated time points. *SCR1* was used as a loading control. **F**, *IRT1* and *IRT2* expression as detected by northern blot of *MAT*a cells harboring *ime1*Δ and the *GAL1-10* promoter integrated - 10 bp from the *Ume6* binding site at *IRT2* site (*pGAL-IRT2*) (FW8759), or the same cells also expressing *GAL4-ER* (FW8758). Samples were taken from cells were grown in YPD till the indicated time-points. *SCR1* was used as a loading control.

In the wake of transcription, nucleosomes disassemble and re-assemble to maintain chromatin structure (Venkatesh and Workman, 2015). We hypothesized that increasing *IRT2* transcription levels promotes chromatin assembly and thus transcriptional repression. At low levels of *IRT2* transcription, the rate of transcription-coupled nucleosome assembly is low, but histone exchange is increased (Figure 5D). Consequently, H3K56ac-containing nucleosomes disassemble to facilitate recruitment of Rme1. Conversely, at higher levels of *IRT2* transcription, nucleosome assembly rates are elevated. Well-positioned nucleosomes, in turn, interfere with Rme1 recruitment and *IRT1* expression. To test this model, we modulated the level of *IRT2* transcription by integrating a range of LexA operator sequence repeats near the *IRT2* transcription start site (lexO_n_(+96 bp from Ume6 binding site)), and measured *IRT2* and *IRT1* levels together with nucleosome positioning (Figure 6B, 6C, S6A, S6B and S6C). Upon activation by LexA-ER with β-estradiol, *IRT2* levels as well as Pol II binding increased with the number of integrated lexO_n_(+96) repeats (Figure 6B, S6A, S6B, and S6D). In accordance with our model, we found that the higher *IRT2* transcription, the greater the repression of *IRT1* transcription. Importantly, a positioned nucleosome was detected encompassing the Rme1 binding sites concomitantly with increasing levels of *IRT2* transcription (Figure 6C and S6C). We conclude that there is transcription dependent effect of *IRT2* on repression of *IRT1*.

Next, we examined whether low levels of *IRT2* transcription induced by the lexO system was sufficient for activating *IRT1* expression. Since we disrupted the Ume6 binding site in lexO_n_(+96) cells, causing low levels of *IRT2* transcription, *IRT1* was induced and no positioned nucleosome was detected near the Rme1 binding sites (Figure S6C). Therefore, we integrated a single lexO site (lexO(−10)) upstream of the Ume6 binding site in *ime1*Δ cells, thus preserving *IRT2* repression (Figure 2F). Remarkably, we were able to rescue the *IRT1* expression defects of *ime1*Δ cells with a single lexO(−10) site without the need to activate LexA-ER with β-estradiol (Figure 6D (see lanes: 17-24)). These cells displayed constitutive low levels of *IRT2* and marked levels of *IRT1* expression across all time points, despite the presence of the Ume6 binding site. The effects on *IRT1* expression by the single lexO(−10) site were partially suppressed by *rtt109*Δ further supporting the role of H3K56ac in *IRT2* mediated activation of *IRT1* (Figure 6E). A similar result was obtained when *IRT2* upstream sequence was under the control of *GAL1-10* promoter (*pGAL-IRT2*) in cells harbouring Gal4-ER (no β-estradiol) (Figure 6F). In the absence of Gal4-ER, we detected neither *IRT2* nor *IRT1* expression in *pGAL-IRT2* cells indicating that Gal4-ER was required for inducing *IRT2* transcription and consequently, *IRT1*. We conclude that low levels of *IRT2* transcription are sufficient to induce *IRT1* expression through transcription coupled histone exchange and nucleosome disassembly, while higher levels of *IRT2* transcription direct nucleosome assembly and thereby repression of *IRT1* expression.

### Mathematic model describing how *IRT2* transcription levels control meiotic entry

The data above demonstrate that low levels of *IRT2* transcription activates *IRT1* expression, while higher levels of *IRT2* transcription repress *IRT1* expression in a dose dependent manner. The levels of *IRT1* transcription are also tightly linked to the levels of Rme1 in the cell, which vary greatly between cells expressing a single mating-type (*MAT*a or *MAT*α) and cells expressing opposite mating-types (*MAT*a/α), and between *MAT*a/α cells of different genetic backgrounds (Deutschbauer and Davis, 2005; Gerke et al., 2009; Mitchell and Herskowitz, 1986). To quantitatively dissect how the different signals of *IRT2* and Rme1 impinge on Ime1 expression, we developed a mathematical model describing the regulatory circuit consisting of *IRT2*, Rme1, *IRT1*, and *IME1* (Figure 7A and Figure S7A). To test the model, we first simulated high and low Rme1 levels representing the single mating-type (*MAT*a or *MAT*α, haploid) and, haploid *MAT*a/α (diploid) cell-type, respectively, which resulted in repression or activation of Ime1 expression (Figure 7B and 7C). Moreover, agreeing with our experimental data, the model predicted that in the absence of *IRT2*, activation of *IRT1* transcription independent of mating-type status, and consequently induced Ime1 expression. To further dissect how Rme1 levels and the dual function of *IRT2* control Ime1 expression, we assessed the dose-response association between Rme1 and Ime1 (Figure 7D, S7B and S7C). The analyses revealed a sigmoidal relationship, where over a wide range of high to low Rme1 levels, Ime1 expression was either repressed or activated. The two extremes of the curve, Ime1 expression or repression, represent the ability for diploid (*MAT*a/α) and inability for haploid (*MAT*a or *MAT*α) cells to enter meiosis. Importantly, the absence of *IRT2* transcription or the presence of only activating low levels of *IRT2* abrogated the bimodal relationship between Rme1 and Ime1 expression levels, supporting the two-sided function of *IRT2* as a requisite for the cell-type specific control of yeast gametogenesis (Figure 7D and Figure S7C).

**Figure 7.**
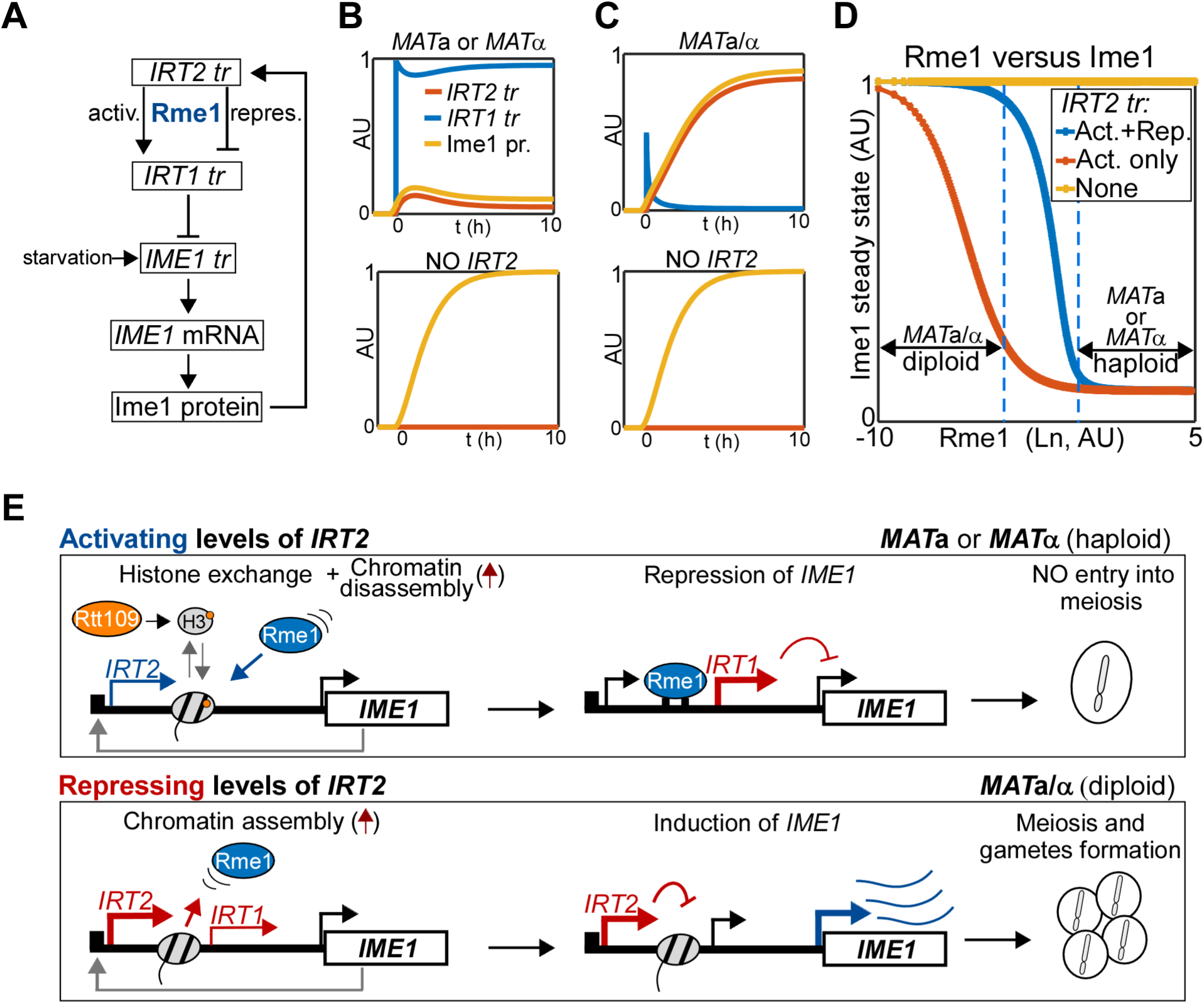
Modelling of mating-type control of Ime1 expression involving two lncRNAs. **A**, Schematic of the mathematical model displaying the regulatory circuit controlling Ime1 expression. **B and C**, Simulation of *MAT*a and *MAT*a/α cells in the presence or absence of *IRT2*. **D**, Simulation of Ime1 steady state protein levels over a range of Rme1 concentrations (Ln) in the presence of either WT levels of *IRT2* (activating and repressive effect on *IRT1*), activating levels *IRT2* (activating effect on *IRT1* only), or no *IRT2*. **E**, Model describing how distinct levels of *IRT2* control the decision of whether or not to enter meiosis and form gametes in haploid (*MAT*a or *MAT*α) and diploid (*MAT*a/α) yeast cells.

## Discussion

Multiple signalling cues act in hierarchical order to control the decision to enter meiosis in yeast. In this work, we demonstrated that distinct transcription levels of the lncRNA *IRT2* ensures a robust transition from nutrient to mating-type control of *IME1* promoter activity. At relative low transcription levels, *IRT2* promotes transcription of the adjacent lncRNA *IRT1*, which in turn represses the *IME1* promoter. Increasing levels of *IRT2* interferes with transcription of *IRT1*, allowing for activation of the *IME1* promoter. The dual role of *IRT2* shapes this regulatory circuit that is set in motion by Ime1 itself, which enables cells expressing opposite mating-type loci (typically diploid cells), but not cells expressing a single mating-type locus (typically haploid cells), to enter meiosis. Our data show that transcription levels of an lncRNA can have a critical role in regulating gene expression and cell fate outcomes.

### Mechanism of *IRT2* mediated activation of transcription

Two lncRNAs, *IRT1* and *IRT2*, are transcribed through different parts of the *IME1* promoter to control *IME1* expression, and thereby the decision to enter meiosis (Moretto et al., 2018; van Werven et al., 2012). Several lines of evidence indicate that *IRT2* directly promotes *IRT1* transcription in cells with a single mating-type. First, partial deletions in *IRT2*, without affecting the Rme1 binding sites, compromises Rme1 recruitment and activation of *IRT1* transcription. Second, insertion of a transcriptional terminator between the Rme1 binding sites and the *IRT2* transcription start site affected *IRT1* activation, suggesting that the act of transcription is required. Third, altered *IRT2* transcription patterns due to mutations in Ime1 or the Ume6 binding sites upstream in the *IME1* promoter, affected *IRT1* expression in a comparable way. Finally, low levels of *IRT2* transcription controlled from a heterologous promoter directly upstream of *IRT2* was sufficient to induce *IRT1* transcription.

How does *IRT2* promote *IRT1* activation? In a small-scale gene deletion screen, we identified Rtt109 as a regulator of *IRT1* transcription. As a result, cells with a single mating-type can enter meiosis when *RTT109* is deleted or when the main substrate of Rtt109, H3K56ac, is directly disrupted. We propose that low levels of *IRT2* transcription stimulates H3K56ac incorporation into nucleosomes locally. The H3K56ac mark, in turn, facilitates Rme1 recruitment, and activation of *IRT1* transcription. Our results are consistent with a model describing that H3K56ac increases nucleosome unwrapping facilitating transcription factor (TF) binding and transcription activation (Bernier et al., 2015; Neumann et al., 2009; Williams et al., 2008). At *IRT2*, partial unwinding of H3K56ac nucleosomes may provide access for Rme1 binding and thus subsequent *IRT1* activation.

Our data suggest that *IRT2* transcription stimulates nucleosome turnover, possibly directing H3K56ac to chromatin locally. However, how exactly *IRT2* transcription stimulates H3K56ac incorporation into chromatin remains to be determined. One possibility is that chaperones or other factors travel along with Pol II to facilitate incorporation of acetylated histones into nucleosomes (Park and Luger, 2008). During DNA replication in yeast, H3K56ac is directed to nucleosomes by chromatin assembly factors, perhaps they also play a role at *IRT2* (Kaplan et al., 2008; Topal et al., 2019). Another possibility is that in the wake of transcription partial disassembly of nucleosomes leads to stochastic exchange of histones (Jamai et al., 2007). In line with this idea, H3K56ac is enriched in transcribed gene bodies in the *set2* deletion mutant, thus in the absence of H3K36 methylation, suggesting that transcription can promote histone exchange (Venkatesh et al., 2012). Once the H3K56ac is incorporated into nucleosomes, the mark itself may promote nucleosome turnover (Kaplan et al., 2008; Rufiange et al., 2007). In this regard, *IRT2* transcription is perhaps the first event that directs H3K56ac to chromatin, subsequently H3K56ac can act in positive feedback to further increase nucleosome turnover. It is worth noting that H3K56ac is widespread at promoter proximal nucleosomes where divergent noncoding transcription takes place, which raises the interesting possibility that H3k56ac incorporation via noncoding transcription may be widespread (Topal et al., 2019; Xu et al., 2009).

The transcription-controlled histone exchange and incorporation of acetylated histones into chromatin as we described here, could be reminiscent of how a class of lncRNAs called enhancer RNAs (eRNAs) regulate gene expression (Li et al., 2016). Like *IRT2*, eRNAs are typically very lowly expressed, facilitate recruitment of transcription factors, and the act of eRNA transcription has been shown to be important for enhancer activity. Production of eRNAs promotes enhancer activity and activation gene expression. Expression of eRNAs also correlates with certain histone marks including histone acetylation. Interestingly, in mammalian cells H3K56ac is enriched at active enhancers and promoters, suggesting that noncoding transcription directed H3K56ac could be conserved (Skalska et al., 2015; Tan et al., 2013).

### Model for the dual function of transcription of an lncRNA

Previously, we described the regulatory circuit consisting of *IRT2*, *IRT1*, and *IME1* (Moretto et al., 2018). We showed that *IRT2* transcription interferes with *IRT1*, which in turn leads to up-regulation of *IME1* expression and entry into meiosis in *MAT*a/α cells. In this work, we demonstrated *IRT2* activates *IRT1* transcription in cells with a single mating-type. Moreover, we showed that *IRT2* transcription levels play a determining role. The *IRT2* effect on gene regulation follows a hormetic pattern. In absence of *IRT2* transcription, no activation of *IRT1* transcription will occur. However, relatively low levels of *IRT2* will promote *IRT1* activation, while increasing *IRT2* transcription will repress *IRT1* (Figure 7E).

How do transcription levels of *IRT2* determine whether to activate or repress gene expression? Our data suggest that there is a dynamic interplay between nucleosome assembly and disassembly, and transcription factor concentration. In the wake of transcription, nucleosomes disassemble and re-assemble to maintain chromatin structure (Ard et al., 2017; Venkatesh and Workman, 2015). With this view, as the transcription levels of *IRT2* increases the rate of transcription-coupled nucleosome assembly also increases, eventually leading to repression of *IRT1*. In context activation, *IRT2* transcription directs histone H3K56ac to nucleosomes, which in turn facilitates nucleosome disassembly. In principle, one round of transcription can be sufficient to direct histone exchange, and thus nucleosome disassembly. When the rate of transcription coupled nucleosome assembly is higher than the rate of nucleosome disassembly, then the “tip the scale” toward repression will occur (Figure 7E). As depicted in our mathematical model, the concentration of Rme1 also plays an important role in activation of *IRT1* transcription. The higher Rme1 levels are in the cell, the earlier a stable association to its binding site will occur, and thus an activation of *IRT1* transcription. Our model further shows that the dual function of *IRT2* and Rme1 concentration in the cell form a circuit to regulate mating-type signalling to Ime1 expression. Taken together, our study of the *IME1* promoter demonstrates that noncoding transcription levels can play a determining role on whether repression or activation of gene transcription will occur, thereby controlling gene expression outcomes.

Many examples of gene regulation by lncRNA transcription with a range of different outcomes have been reported (Engreitz et al., 2016; Gil and Ulitsky, 2019; Hirota et al., 2008; Kornienko et al., 2013; Martens et al., 2004; van Werven et al., 2012). Our finding that distinct transcription levels of an lncRNA direct opposing chromatin and transcription states illustrates the gene regulatory potential for transcription of lncRNAs in general. Given that lncRNAs are typically expressed across many parts of the genome, from yeast to humans, we propose that the act of transcription itself through promoters or other regulatory regions may have extensive functions in determining gene expression levels (David et al., 2006; Hon et al., 2017; Iyer et al., 2015; Kung et al., 2013).

## Supplementary Information

Methods, References, Figures S1-S7 legends, Figures S1-S7 and Table S1-S4

## Author Contributions

FM and FW conceived and designed the study; FM and FW designed the experiments; FM performed the experiments; FM and FW analyzed the data; EW generated mathematical model; MC, CL, and NL helped with generating and analyzing sequencing data; FM and FW wrote the manuscript with input from all authors. FW supervised the project.

## Declaration of Interests

The authors declare no competing interests.

## Acknowledgements

We are grateful to Frank Uhlmann, Jesper Svejstrup and members of the Van Werven lab for advice and critical reading of the manuscript. We thank Elçin Ünal, Philipp Korber, Valerie Borde, and Celine Bouchoux for sharing reagents. NEW is supported by Cancer Prevention and Research Institute of Texas (RR150058), which are obtained by Andreas Doncic. FM and FW are supported by the Francis Crick Institute (FC001203), which receives its core funding from Cancer Research UK (FC001203), the UK Medical Research Council (FC001203), and the Wellcome Trust (FC001203). CL and NL are supported by the Francis Crick Institute (FC010110), which receives its core funding from Cancer Research UK (FC010110), the UK Medical Research Council (FC001203), and the Wellcome Trust (FC010110). M.C. is supported by a fellowship from the Agency for Science, Technology and Research (A∗STAR) of Singapore.

