## Supplementery information for "Transcription levels of a long noncoding RNA orchestrate opposing regulatory and cell fate outcomes in yeast"

**Supplementary Information**

Methods

References

Figures S1-S7 legends

Figures S1-S7

Tables S1-S4

**METHODS:**

**Yeast strains and plasmids**

Yeast strains used in this paper were derived from the SK1 strain background. Gene or promoter deletions were generated using the one-step deletion protocol as described previously (Longtine et al., 1998). The *RPB3-AID* allele was generated through a one-step C-terminal tagging procedure of *RPB3* with an auxin-inducible degron (AID) tagging cassette, which contains the IAA17 degron and three copies of the V5 epitope. The *RPB3-FLAG* allele was generated through a one-step C-terminal tagging procedure of *RPB3* with a FLAG tag cassette which contains three copies of the FLAG epitope (gift Jesper Svejstrup). The *RPB3-AID* strain also expressed the *Oryza sativa TIR1* (*osTIR1*) ubiquitin E3 ligase from the *CUP1* promoter (pNH603-pCUP1-osTIR1, gift from Elçin Ünal). The pNH603-pCUP1-osTIR1 plasmid was linearized by digestion with PmeI and integrated as a single copy at the *HIS3* locus. *MAT*a/a diploid strains were generated by replacing the *MAT*α locus in *MAT*a/α diploids with a plasmid linearized by EcoRI digestion harboring the *MAT*a fragment with *URA3* or *TRP1* selectable markers (pRS304-*MAT*a, this work; pRS306-*MAT*a)(van Werven et al., 2012). The *irt2-T* allele harbors the *CYC1* terminator sequence, which was integrated in the *IRT2* locus using the ‘‘delitto perfetto’’ strategy (Storici et al., 2001). In short, *Kluyveromyces lactis URA3* marker was first integrated in the *IRT2* locus, subsequently a PCR product containing the *CYC1* terminator and homology to the flanking sequences was used to excise the *URA3* marker by 5-fluoro-orotic acid counter selection. As a control we also generated an *irt2-I* allele, which harbors a control insert from pUG6-Myc-Avitag (van Werven and Timmers, 2006). To excise the *KanMX* marker from *irt2-I*, we expressed the Cre recombinase from a plasmid (pRS304-GPDpr-CRE-EBD78-CYC1t, which is pTW040 re-cloned into pRS304, gift from Celine Bouchoux)(Terweij et al., 2013). After excision of the *KanMX* marker, genetic crosses were used to remove the pRS304-GPDpr-CRE-EBD78-CYC1t from *irt2-I*. Histone H3K56A and H3K56R mutants were generated through site directed mutagenesis using a plasmid carrying histone H3 and H4 genes along with a *URA3* marker (*HHT1* and *HHF1*,pDM9 (Sommermeyer et al., 2013), gift from Valerie Borde). Mutant plasmids were transformed into a strain with all four genomic copies of histone H3 and H4 genes deleted and harboring a plasmid wild-type for the histone H3 and H4 genes on TRP1 selectable marker (*HHT2* and *HHF2,* pVB140 (Sommermeyer et al., 2013), gift from Valerie Borde). Genetic crosses were used to generate the histone H3K56A and H3K56R mutants in the absence of the wild-type histone H3 covering plasmid. The plasmids for generating strains with LexA operator (lexO) sequences and LexA fused to the estrogen receptor domain and activation domain (LexA-ER-HA-B112) were described previously (gift from Elçin Ünal) (Chia et al., 2017; Ottoz et al., 2014).  A different number of LexO sites were integrated at *IRT2* locus position +96 or -10 bp from the Ume6 binding site. A plasmid expressing LexA-ER-HA-B112 from the *GPD1* promoter was linearized with SfiI restriction enzyme and integrated at the *TRP1* locus. The strains used for the histone H3 exchange assay described in Figure 5C and S5 were described previously (Schermer et al., 2005). In short, we used a strain with the endogenous copies of histone H3 and H4 deleted and covered by a centromeric plasmid pNOY439 harboring *HHT2*-Myc (histone Myc-H3) and untagged *HHF2* (histone H4) under control of their endogenous promoters. In addition this stain also harbored a plasmid (YIplac211 pGAL1-10 HHF1 FLAG-HHT1) integrated at the *URA3* locus with *HHT1* FLAG tagged (histone FLAG-H3) and untagged *HHF1* (histone H4) under control of the *GAL1-10* promoter (Schermer et al., 2005). To generate *pGAL-IRT2* strain, the

*GAL1-10* promoter was integrated 10 bp upstream of the Ume6 binding site. The *GAL4-ER* expression construct was described previously (Carlile and Amon, 2008). The strain genotypes and plasmids are listed in Table S2 and S4.

**Growth and conditions**

All experiments were performed at 30°C in a shaker incubator at 300rpm. A protocol for rapid induction of *IME1* or *IRT1* expression was described previously (van Werven et al., 2012). In short, cells were grown till saturation for 24h in YPD (1.0% (w/v) yeast extract, 2.0% (w/v) peptone, 2.0% (w/v) glucose, and supplemented with uracil (2.5 mg/l) and adenine (1.25 mg/l)), cells were then diluted at $OD_{600}$ = 0.4 to pre-sporulation medium (1.0% (w/v) yeast extract, 2.0% (w/v) bacto tryptone, 1.0% (w/v) potassium acetate, 50 mM potassium phthalate) grown for about 16 hours, subsequently centrifuged, washed with sterile miliQ water, centrifuged again, re-suspended at $OD_{600}$ = 1.8 in sporulation medium (SPO) (0.3% (w/v) potassium acetate and 0.02% (w/v) raffinose)) and incubated at 30°C. For Figures 1C, 2F, 2H and S2G, we used a protocol inducing *IRT1* or *IME1* expression with slow kinetics. In short, cells were grown till saturation for 24h in YPD then diluted in YPD aiming to reach $OD_{600}$ = 6 after 16h, several samples were taken from that point till 24h, subsequently cells were washed, and transferred to SPO.

For the RNA polymerase II depletion experiment using Rpb3-AID described in Figure 2D, S2D, and S2E, cells were grown till saturation for 24h in YPD, diluted, and grown for another 16h till OD600 = 6, subsequently cells were split and treated with auxin 500 μM or DMSO for 4h. $CuSO_4$ (50μM) was also added to allow expression of the

Tir1 ligase from *pCUP1-osTIR1* integrated at the *HIS3* locus. Cells were then washed, transferred to SPO at OD600 = 1.8 and incubated at 30°C for 3h before ChIP and protein samples were collected.

For the lexO/LexA-ER experiments described in Figure 6B-E and S6A-C cells were grown till saturation for 24h in YPD, diluted ($OD_{600}$ = 0.4) in pre-sporulation medium and grown for another 16h. Cultures were then split and treated with $\beta$-estradiol (25nM) or ethanol (mock) for 1h, transferred to SPO ($OD_{600}$ = 1.8), and incubated up till 3h in the presence of $\beta$-estradiol (15 nM) or ethanol.

For the and *pGAL-IRT2*/GAL4-ER experiment described in Figure 6F, cells were till saturation for 24h in YPD, diluted ($OD_{600}$ = 0.4) in pre-sporulation medium and grown for another 16h. Subsequently samples were taken for the indicated time points.

For measuring histone exchange rates described in Figure 5 and S5, cells were grown till saturation for 24h in YPD then shifted and grown overnight in YP raffinose 2% (YPR) till it reached $OD_{600}$=0.9. Cells were then arrested in G1 with $\alpha$ factor (5 $\mu$g/ml) for 2h, subsequently split to YP plus 2% galactose and YPR both containing $\alpha$ factor (5 $\mu$g/ml).

**Nuclei/DAPI counting**

DAPI staining was used to monitor meiotic divisions throughout time courses. Cells were fixed in 80% (v/v) ethanol, pelleted by centrifugation and re-suspended in 100 mM phosphate buffer (pH 7) with 1 μg/ml 4′,6-diamidino-2-phenylindole (DAPI). Cells were then sonicated for a few seconds and left in the dark at room temperature for at least 5 minutes. The proportion of cells containing one, two, three or four (meiosis I + II) DAPI masses were counted using a fluorescence microscope.

**Chromatin immunoprecipitation**

Chromatin immunoprecipitation (ChIP) experiments were performed as described previously (Moretto et al., 2018). Cells were fixed in 1.0% w/v formaldehyde for 25 minutes at room temperature and quenched with 100 mM glycine. Cells were lysed in FA lysis buffer (50 mM HEPES–KOH, pH 7.5, 150 mM NaCl, 1mM EDTA, 1% Triton X-100, 0.1% Na-deoxycholate, 0.1% SDS and protease cocktail inhibitor used as recommended by the manufacturer (complete mini EDTA-free, Roche)) using beadbeater (BioSpec) and chromatin was sheared by sonication using a Bioruptor (Diagenode, 8 cycles of 30 seconds on/off). Extracts were incubated for 2 hours at room temperature with anti-V5 agarose beads (Sigma) or overnight at 4°C with magnetic Prot A beads (Sigma) coupled with a polyclonal antibody raised against Histone H3 (Ab1791, Abcam) or Histone H3 acetylated lysine 56 (07-677-I, Millipore), washed twice with FA lysis buffer, twice with wash buffer 1 (FA lysis buffer containing 0.5M NaCl), and twice with wash buffer 2 (10 mM Tris–HCl, pH 8.0, 0.25M LiCl, 1 mM EDTA, 0.5% NP-40, 0.5% Na-deoxycholate). Subsequently, reverse cross-linking was done in 1% SDS-TE buffer (100 mM Tris pH 8.0, 10 mM EDTA, 1.0% v/v SDS) at 65°C overnight. After 2 hours of proteinase K treatment, samples were purified, and DNA fragments were quantified by real-time PCR using SYBR green mix (Life Technologies) using primers described in Table S3. Signals were normalized over the *HMR* locus, which showed no binding for Rme1. For Figure 4G, H3K56ac ChIP signals were normalized to histone H3 ChIP. For the histone H3 turnover experiments described in Figure 5C-F and S5D, ChIPs were performed as described above using antibodies against the Myc epitope (clone 9E11, Thermofisher) or against the FLAG epitope (M2 beads, Sigma). Signals were normalized using primers directed against a telomeric region from chromosome VI. For the Rpb3-FLAG ChIPs, we used FLAG-antibody coupled beads (M2 beads, Sigma). Signals were normalized over the *HMR* locus, which showed no binding for Rpb3.

**Micrococcal nuclease (MNase) qPCR**

Nucleosome positioning at the *IRT2* locus was determined by quantifying the abundance of mononucleosomal DNA using a MNase digestion protocol that was described previously (Rando, 2010). In short, approximately 90 $OD_{600}$ units of cells were crosslinked for 25 min at 30°C with 1% (v/v) formaldehyde. Reaction was quenched with the addition of glycine to 125 mM. Subsequently, cells were re-suspended in 20 ml of buffer Z (1 M sorbitol, 50 mM Tris-HCl pH 7.4**)** plus β-mercaptoethanol (10mM) and treated with 250 µg of T100 Zymolyase for 60 min. Next, cells were re-suspended in 1 ml NP buffer (0.5 mM spermidine, 1 mM β-mercaptoethanol (β -ME), 0.075% (w/v) Tergitol solution-type NP-40 detergent (NP-40), 50 mM NaCl, 10 mM Tris-HCl pH 7.4, 5 mM $MgCl_2$ , 1 mM $CaCl_2$), vortexed for 10 seconds, and 100 µl of extract was treated with 0.2 µl of MNase (2mg/ml, NEB) for 30 min at 37°C, the reaction was quenched with 10 mM EDTA, and reverse crosslinked overnight in 1% SDS-TE and 4 units per ml of proteinase K (NEB). Samples were treated with RNase A and purified DNA fragments were separated by gel electrophoresis before gel purification of the mono-nucleosome bands. MNase treated and input samples were quantified by qPCR on a 7500 FAST Real-Time PCR machine (Life Technologies) using Platinum SYBR mix (Thermofisher). The signals were normalized using primers directed against a telomere locus. The scanning primer pairs covering the *IRT2* locus and downstream region used for the analysis are available in Table S3.

##### Northern blotting

Northern blots were performed as described previously (Moretto et al., 2018). In short, total RNA was extracted with Acid Phenol:chloroform:Isoamyl alcohol (125:24:1) and precipitated in ethanol with 0.3 M sodium acetate. RNA samples were denatured in a glyoxal/DMSO mix (1 M deionized glyoxal, 50% v/v DMSO, 10 mM NaPi buffer pH 6.5-6.8) at 70°C for 10 minutes. Samples were mixed with loading buffer (10% v/v glycerol, 2 mM NaPi buffer pH 6.8, 0.4% w/v bromophenol blue) and separated on an agarose gel (1.1% w/v agarose, 0.01 M NaPi buffer pH 6.8) for 2 hours at 80 V. RNAs were then transferred onto nylon membranes overnight by capillary transfer in 0.025 M NaPi buffer pH 6.8. Membranes were blocked for 2-3 hours at 42°C in Hybridization buffer (1% w/v SDS, 50% v/v de-ionized formamide, 25% w/v dextran sulfate, 58 g/L NaCl, 200 mg/L herring sperm single strand DNA, 2 g/L BSA, 2 g/L polyvinyl-pyrrolidone, 2 g/L ficoll, 1.7 g/L pyrophosphate, 50 mM Tris pH 7.5) before hybridization. Radioactive probes were synthesized using a Prime-It II

Random Primer Labeling Kit (Agilent), a target-specific DNA template and dATP [α-32P] (Perkin-Elmer). To avoid differences in signal intensity due to labelling quality and blotting variations, for each experiment samples were run on a single gel, transferred onto a single membrane, and hybridized with probes in a single hybridization tube. Background normalised quantifications of the northern blots were done using Imagej software (Schneider et al., 2012). The oligo nucleotide sequences used to generate target-specific DNA template for amplifying the northern blot probes are displayed in Table S3. We have included a supplementary file with all the raw blots.

**RNA-seq**

At least 5 µg of total RNA was treated with DNAse and purified on column (Macherey-Nagel). At least 500 ng of purified total RNA was used as input for the KAPA mRNA Hyper Prep kit (KK8580, Roche). Libraries were prepared according to manufacturer's instructions. After bead based clean up, libraries were sequenced on an Illumina HiSeq 2500 to an equivalent of 75 bases single-end reads, at a depth of approximately 20 million reads per library.

**TSS-seq and TES-seq**

The TSS sequencing approach was adapted and modified from previously published protocols (Adjalley et al., 2016; Arribere and Gilbert, 2013; Malabat et al., 2015).. At least 5 µg of mRNAs were purified from total RNA using the Poly(A)Purist MAG kit (AM1922, Ambion). mRNAs were fragmented for 3 minutes at 70°C using a Zinc-based alkaline fragmentation reagent (AM8740, Ambion). RNAs were cleaned up using RNeasy MinElute Cleanup Kits (74204, Qiagen) to enrich for 200-300 nt fragments. These fragments were dephosphorylated with 30 units of recombinant shrimp alkaline phosphatase (M0371, NEB) for 1 hour at 37°C with RNasin Plus, the phosphatase was heat inactivated and the RNA was extracted with Acid Phenol:chloroform: Isoamyl alcohol (125:24:1) and precipitated at -20°C overnight in ethanol with 0.3M sodium acetate and 1 µl linear acrylamide (AM9520, Ambion). RNA was then subjected to a decapping reaction with 2 units of Cap-Clip acid pyrophosphatase (C-CC15011H, Tebu-Bio) and with RNasin Plus. RNAs were then extracted using acid Phenol:chloroform:isoamyl alcohol (125:24:1) and precipitated in ethanol. Some RNA from a SPO (starvation) 0 hour time point was set apart without the decapping reaction as a non-decapping control. Subsequently, the RNA was mixed with 10 µM of custom 5' adapter (CACTCTrGrArGrCrArArUrArCrC) and the ligation reaction was done using T4 RNA ligase 1 (M0437M, NEB) and with RNasin Plus. The ligation reaction was cleaned up with the RNeasy MinElute Cleanup Kit and RNAs were mixed with 2.5 µM random hexamers (N8080127, ThermoFisher Scientific) and RNasin Plus, denatured at 65°C for 5 minutes and cooled on ice. Reverse transcription reactions were carried out using SuperScript IV reverse transcriptase (18090010, Invitrogen) at 23°C for 10 minutes, 50°C for 10 minutes, 80°C for 10 minutes and held at 4°C. The RNA templates were degraded by incubating reactions with 5 units of RNase H (M0297, NEB) and 1.0 µl of RNase cocktail enzyme mix (AM2286, Ambion). DNA products were purified using 1.8x volume of HighPrep PCR beads (AC-60050, MagBio). Purified products were subjected to second strand synthesis using 0.3 µM of second strand biotinylated primer (GCAC/iBiodT/GCACTCTGAGCAATACC) and the KAPA Hi-Fi hot start ready mix (KK2601, Roche). The second strand reaction was carried out at 95°C for 3 minutes, 98°C for 15 seconds, 50°C for 2 minutes, 65°C for 15 minutes and held at 4°C. Double stranded product (dsDNA) was purified with 1.8x volume HighPrep PCR beads and concentration was quantified using the Qubit dsDNA HS assay kit (Q32851, Invitrogen). At least 1 ng of dsDNA was then used as input for the KAPA Hyper Prep Kit (KK8504, Roche) and ligated to KAPA single indexed adapters Set A (KK8701, Roche) or Set B (KK8702, Roche). Samples were processed according to manufacturer's instructions with one exception: just prior to the library amplification step, samples were bound to MyOne Streptavidin C1 Dynabeads (65001, ThermoFisher Scientific) to capture biotinylated dsDNA. Library amplification was done on the biotinylated dsDNA fraction bound to the beads. Depending on the input amounts, 15 PCR cycles were used to generate libraries. Amplified libraries were quantified by Qubit, and adapter-dimers were removed by electrophoresing libraries on Novex 6% TBE gels (EC62655BOX, Invitrogen) at 120 V for 1 hour, and excising the smear above ~150 bp. Gel slices containing libraries were shredded by centrifugation at 13000 g for 3 minutes. Gel shreds were re-suspended in 500 μl crush and soak buffer (500 mM NaCl, 1.0 mM EDTA and 0.05% v/v SDS) and incubated at 65°C for 2 hours on a thermomixer (1400 rpm for 15 seconds, rest for 45 seconds). Subsequently, the buffer was transferred into a Costar SpinX column (8161, Corning Incorporrated) with two 1 cm glass pre-filters (1823010, Whatman). Columns were centrifuged at 13000 g for 1 minute. DNA libraries in the flowthrough were precipitated at -20°C overnight in ethanol with 0.3 M sodium acetate and 1 μl linear acrylamide (AM9520, Ambion). Purified libraries were further quantified and inspected on a Tapestation (Agilent Technologies) and sequenced on an Illumina HiSeq 2500 to an equivalent of 75 bases single-end reads, at a depth of approximately 20 million reads per library.

The TES sequencing approach was adapted and modified from previously published protocols (Lai et al., 2015; Ng et al., 2005). From the same pool of fragmented mRNAs describe above in the 5' end sequencing, at least 1 µg was used for 3' end sequencing. RNA fragments were mixed with 2.5 µM GsuI20TVN primer (/5BiotinTEG/ GAGCTAGTTCTGGAGTTTTTTTTTTTTTTTTTTTTVN), 0.5 mM 5-Methylcytosine-dNTPs (D1030, Zymo Research) and 0.5 µl RNasin Plus. Reaction mixtures were denatured at 65°C for 5 minutes and held at 50°C without allowing to cool. SuperScript IV, reaction buffer and 0.4 µg of Actinomycin D were added to the hot reaction mixtures and reverse transcription was performed at 50°C for 10 minutes, 80°C for 10 minutes and held at 4°C. Samples were cleaned with 1.8x volume HighPrep beads and biotinylated RNA:DNA hybrids were captured on MyOne Streptavidin C1 Dynabeads. After capture, streptavidin beads were washed once with 1x NEbuffer 2 (B7002S, NEB), re-suspended in water and subjected to second strand synthesis. The 50 µl second strand synthesis reaction consisted of 20 µl re-suspended streptavidin beads, 1X NEbuffer 2, 250 µM dNTPs, 26 µM NAD+ (B9007S), 2.5 units RNase H, 10 units E.coli DNA ligase (M0205S), and 15 units DNA polymerase I (M0209S). Second strand synthesis reactions were conducted at 16°C for 2.5 hours on a thermomixer (1400 rpm for 15 seconds, rest for 2 minutes). After reaction, beads were washed once with 1x binding and washing buffer (5.0 mM Tris-HCl pH 7.5, 0.5 mM EDTA, 1.0 M NaCl) and once with buffer B (10 mM Tris-HCl pH 7.5, 10 mM MgCl2, 0.1 mg/ml BSA). Washed beads were re-suspended in 18 µl buffer B and digested with 10 units of GsuI (ER0461, ThermoFisher Scientific) at 30°C for 1 hour on a thermomixer (1400 rpm for 15 seconds, rest for 2 minutes). After digestion, the DNA fragments in the supernatant were extracted with

Phenol/chloroform and precipitated at -20°C overnight in ethanol with 0.3 M sodium acetate and 1 μl linear acrylamide. The concentration was quantified using the Qubit dsDNA HS assay kit. At least 1 ng of dsDNA was then used as input for the KAPA Hyper Prep Kit (KK8504, Roche) and ligated to KAPA single indexed adapters Set A (KK8701, Roche) or Set B (KK8702, Roche). Samples were processed according to manufacturer's instructions. Amplified libraries were cleaned and purified by gel extraction using the procedures described in the previous section for TSS sequencing. Purified libraries were further quantified and inspected on a Tapestation (Agilent Technologies) and sequenced on an Illumina HiSeq 2500 to an equivalent of 75 bases single-end reads, at a depth of approximately 20 million reads per library.

**Nascent RNA-Seq**

The nascent RNA sequencing procedure was adapted as described in (Churchman and Weissman, 2011). Briefly, about one liter of yeast culture was spin down at the harvest time point and pellet snap frozen in liquid nitrogen. Cell pellets were then grinded in fine powder using a cryo-mill instrument with the instrument maintained cooled with liquid nitrogen. All the powder was re-suspended in about 12 to 15 ml of lysis buffer (20 mM HEPES pH 7.4, 110 mM K acetate, 0.5% triton X-100, 0.1% Tween-20, 10 mM MnCl2, 1X protease inhibitor cocktail HALT, SUPERasin RNAse inhibitor 50 U/ml) and lysate subjected to DNAse I treatment (1200 U per sample) for 20 min. Lysate was then clarified through two rounds of centrifugation at about 20 000g for 10 min at 4C. Anti-FLAG immunoprecipitation was performed using 750 μL of anti-FLAG M2 agarose beads per sample and for 2.5 h at 4C on a rotating wheel. Beads were then washed 5 times using 10 ml of cold wash buffer (20 mM HEPES pH 7.4, 110 mM K acetate, 0.5% triton X-100, 0.1% Tween-20, 10 mM MnCl2, SUPERasin RNAse inhibitor 1 U/ml, 1 mM EDTA) for 5 min at 4C. Beads were then cleaned using a Pierce column and two rounds of elution were performed using FLAG peptide (450 μl 10 mg/ml Sigma FLAG peptide) for 30 min at 4C. Elution was then subjected to a phenol:Chloroform extraction and RNA precipitated overnight in ethanol containing 0.3 M Na acetate. RNAs were then resuspended in Nuclease free water, quantified by Qbit RNA high sensitivity assay kit (Q32852, ThermoFisher Scientific) and analyzed by Bioanalyser (Agilent technologies).

At least 500 ng of purified RNA was treated with DNAse and purified on column (Macherey-Nagel). Then, about 100 ng of purified total RNA was used as the inputs for the KAPA RNA hyperPrep kit (KK8540, Roche). Libraries were prepared according to manufacturer's instructions. After bead based clean up, libraries were sequenced on an Illumina HiSeq 4000 to an equivalent of 75 bases single-end reads, at a depth of approximately 50 million reads per library.

**Bioinformatic analysis**

For RNA-seq data (including nascent RNA-seq), adapter trimming was performed with cutadapt (version 1.9.1) with parameters "-a AGATCGGAAGAGCACACGTCTGAACTCCAGTCAC --minimum-length=20". STAR (version 2.5.2) with parameters "--alignIntronMin 3 --alignIntronMax 5000" was used to perform the read mapping to the *S. cerevisiae* SK1 genome assembly from Keeney lab (http://cbio.mskcc.org/public/SK1_MvO/) (Martin, 2011). Alignments with mapping quality of <10 or soft/hard-clipping were filtered (Dobin et al., 2013; Martin,

2011). Alignments from forward strand and reverse strand were separated by using "samtools view -b -f 0x10" and "samtools view -b -F 0x10" to split the alignments. The tool “bedtools genomecov” (Quinlan and Hall 2010) was used to generate the RNA-seq BedGraph tracks across the genome, for both forward and reverse strands. All the reads mapping to the rRNA were trimmed from the analyses prior the normalization. The BedGraph tracks were normalized by the number of usable reads in each library. BedGraph files were converted to bigWig using the tool bedGraphToBigWig from UCSC (Kent et al., 2010).

For the TSS transcript sequencing data, the custom 5’ adapter sequence specific to the protocol was removed by re-running cutadapt with the parameters “-g CACTCTGAGCAATACC -O 16 --minimum-length=20”, and only the reads containing the adapter sequence were used for further analysis. STAR (version 2.5.2) with parameters “--alignIntronMin 2 --alignIntronMax 1” (i.e. not allowing introns) was used to align TSS-seq reads to the SK1 genome assembly (plus three spike-in sequences) (Dobin et al., 2013). The alignments with mapping quality of >=10 were kept for further analysis. Alignments from forward strand and reverse strand were separated by using "samtools view -b -f 0x10" and "samtools view -b -F 0x10". The 5’-most nucleotide of aligned reads were extracted to generate the genome-wide tracks of TSSs. The BedGraph tracks were normalized by the number of usable reads in each library.

For the TES transcript sequencing data, Adapter trimming was performed with cutadapt (version 1.9.1) with parameters “-a AGATCGGAAGAGCACACGTCTGAACTCCAGTCAC --minimum-length=20”.

STAR (version 2.5.2) with parameters “--alignIntronMin 2 --alignIntronMax 1” as for the 5’ end sequencing part, and alignments with mapping quality of >=10 were kept for further analysis. The reads kept were those with soft-clipping at the 3’ end (size of soft-clipping part <=10) and with at least two consecutive non-templated as in the soft-clipping part. Insertions/deletions were also not allowed. Alignments from forward strand and reverse strand were separated by using "samtools view -b -f 0x10" and "samtools view -b -F 0x10". The 3’-most nucleotide of aligned reads were extracted to generate the genome-wide tracks of TESs. The BedGraph tracks were normalized by the number of usable reads in each library.

**Single molecule RNA fluorescent in situ hybridization (FISH)**

The single molecule RNA FISH was performed as described previously (Moretto et al., 2018; Raj et al., 2008). In short, cells were fixed with formaldehyde overnight, treated with zymolyase and further fixed in 80% ethanol. Subsequently cells were hybridized with fluorophore labelled probes directed to *IME1* (AF594) and *ACT1* (Cy5) as an internal control. Cells were imaged using a 100x oil objective, NA 1.4, on a Nikon TI-E imaging system (Nikon). DIC, DAPI, AF594 (*IME1*), Cy5 (*ACT1*) images were collected every 0.3 micron (20 stacks) using an ORCA-FLASH 4.0 camera (Hamamatsu) and NIS-element software (Nikon). ImageJ software was used to make maximum intensity Z projections of the images (Schneider et al., 2012). StarSearch software (http://rajlab.seas.upenn.edu/StarSearch/launch.html, Raj laboratory, University of Pennsylvania) was used to quantify transcripts in single cells. Comparable thresholds were used to count RNA foci in single cells. Only cells positive for the internal control *ACT1* were used for *IME1* analysis. At least a total n = 150 cells were counted for each experiment.

##### Western blotting

Western blots were performed as previously described (Moretto et al., 2018). Protein extracts were prepared using the trichloroacetic acid (TCA) extraction protocol. After SDS-polyacrylamide gel electrophoresis (4-20% gradient), proteins were transferred onto PVDF membranes. The membranes were then incubated overnight primary antibodies in blocking buffer. Mouse anti-v5 (R96025, Sigma-Aldrich (MO, USA)) was used at a 1:2000 dilution and rabbit anti-hexokinase antibody (H2035, Stratech (Newmarket, UK)) at a 1:8000 dilution. Membranes were then washed in PBST buffer and incubated with IRDye 800CW goat anti-mouse and IRDye 680RD donkey anti-rabbit secondary antibodies (LI-COR (NE, USA)) at a 1:15000 dilution for LI-COR detection. Protein levels were detected on an Odyssey Imager for LI-COR detection.

##### Spot growth assay

Cells were grown using the protocol for rapid induction of *IRT1*. After 0 or 14 days in SPO cells were diluted to $OD_{600}$ = 1 in sterile water, and serial dilutions (5-fold) were spotted onto YPD agar plates. Cells were incubated at 30 °C for 2 days before imaging.

##### Statistics

Statistical significance and tests are indicated in the figure legends and were performed using GraphPad Prism 7 and 8. When comparing only two groups of samples, we have used a parametric two-tailed Student’s *t*-test. When assessing the variance of more than two groups we performed an ANOVA analysis (one way or two ways in function of the experimental design), followed by a Fisher LSD test. Fisher’s LSD test confer more power to the test, compared to performing multiple Students *t*-tests, in that *t*-tests compute the pooled standard deviations from only the two groups being compared, while the Fisher's LSD test computes the pooled standard deviations from all the groups while assuming that all population have similar standard deviation.

**Mathematical modelling**

To investigate of the complex regulation of *IME1* expression by *IRT1* and *IRT2*, we developed a mathematical model. We incorporated new observations described in this manuscript, into the model described previously (Moretto et al., 2018). Specifically, the new mathematical model describes both the activating and repressing properties of *IRT2* on *IRT1* expression. The *IRT2* activating property is incorporated by controlling the *IRT1* transcription rate with the indicator function, $\chi$, which is 1 when *IRT2* transcription rate reaches an activating level threshold, $\mathrm{A}$. After the activation of *IRT1* $\chi$ remains at 1 even when *IRT2* transcription rate falls below $\mathrm{A}$. In addition, the model includes Rme1 concentration, $r$, which is assumed to be constant during the time-period of Ime1 activation. By changing Rme1 concentrations, $r$, we can simulate both haploid and diploid cell-types, which differ significantly in their Rme1 levels. The model variables are listed below (Table 1). The parameters have been chosen to reproduce the qualitative behavior of this system (Table 2).

**Table 1: List of variables (scaled between 0 and 1)**

| | |
|---|---|
| $s$ | Starvation signal |
| $I1$ | *IRT1* transcription rate |
| $I2$ | *IRT2* transcription rate |
| $I_t$ | Ime1 transcription rate |
| $I_m$ | Ime1 mRNA concentration |
| $I_p$ | Ime1 protein concentration |

**Table 2: List of parameters used for simulation**

| | | |
|---|---|---|
| $k_1$ | controls the strength of $I1$ inhibition by $I2$ | 5 |
| $k_2$ | $I_p$ threshold for activating $I2$ | 0.05 |
| $k_3$ | controls the strength of $I_t$ inhibition by $I1$ | 10 |
| $k_4$ | degradation rate of $I_m$ | 1 /h |
| $k_5$ | synthesis rate of $I_p$ | 1 /h |
| $k_6$ | degradation rate of $I_p$ | 1 /h |
| $r$ | Rme1 concentration | 0-5 |
| $A$ | $I2$ threshold for activating $I1$ | 0.01 |

**Equations**

$$s(t) = 1, \qquad \text{for } t \geq 0 \qquad (1)$$

$$\chi = \begin{cases} 1, & I2 \geq A \text{ or } I1 > 0 \\ 0, & \text{otherwise} \end{cases} \qquad (2)$$

$$I1 = \begin{cases} \chi \dfrac{\mathrm{r}}{\mathrm{r} + \mathrm{k_1 I2}}, & r > 0 \text{ or } I2 > 0 \\ 0, & \text{otherwise} \end{cases} \quad (3)$$

$$I2 = \begin{cases} \mathrm{I_p} - \mathrm{k_2}, & Ip \geq \mathrm{k_2} \\ 0, & \text{otherwise} \end{cases} \quad (4)$$

$$\frac{\mathrm{dI_m}}{\mathrm{dt}} = \begin{cases} \dfrac{\mathrm{s}}{\mathrm{s} + \mathrm{k_3 I1}} - \mathrm{k_4 I_m}, & s > 0 \\ -\mathrm{k_4 I_m}, & \text{otherwise} \end{cases} \quad (5)$$

$$\frac{\mathrm{dI_p}}{\mathrm{dt}} = \mathrm{k_5 I_m} - \mathrm{k_6 I_p} \quad (6)$$

The model is simulated with the initial conditions $\mathrm{I_m}(0) = 0$ and $\mathrm{I_p}(0) = 0$.

The first term in Equation (5), $\frac{\mathrm{s}}{\mathrm{s+k_3 I1}}$, describes the *IME1* transcription rate.

Equation (1) models the starvation signal as a step input. Upon starvation, *IME1* mRNA $\mathrm{I_m}$ and in turn Ime1 protein $\mathrm{I_p}$ are synthesized. The evolution of $\mathrm{I_m}$ is given by Equation (5) and of $\mathrm{I_p}$ is given by Equation (6). When $\mathrm{I_p}$ reaches the threshold $\mathrm{k_2}$, *IRT2* transcription, $I2$, turns on. Here we assumed that *IRT2* transcription closely follows Ime1 protein concentration (Equation (4)). Once $I2$ reaches the threshold $A$, the indicator function χ is set at 1 (Equation (2)) and *IRT1* transcription starts (Equation (3)). Note that χ remains equal to 1, even if $I2$ falls below $A$, as long as $I1$ is greater than 0. $I1$ is driven by the Rme1 concentration, $r$, which is assumed to be constant in the cell through the time period of Ime1 activation. Activation of $I1$ suppresses $Ip$ to various degrees depending on $r$ (Figure 7A and S7A). The mathematical model shows how *IRT1* and *IRT2* control expression of *IME1* and Ime1 protein. Activating *IRT2* and no *IRT2* transcription are simulated by setting $I2 = A$

and $I2 = 0$, respectively, for $t > 0$ (Figure 7D and S7C). The mathematical model is simulated in MATLAB 2018a using ode45 function. Code is available upon request.

**Figure S1-S7 legends:**

**Figure S1. *IRT2* is required for *IRT1* expression**

**A,** Schematic of growth conditions used throughout this study. Rapid induction of *IRT1* or *IME1* expression was achieved by growing cells in rich medium (YPD) till saturation (24 h), subsequently cells were shifted to pre-sporulation (pre-SPO) medium and grown for another 16 h. After a wash with sterile water, cells were transferred into sporulation medium (SPO). Slow induction of *IRT1* or *IME1* expression was achieved by shifting cells directly after growth till saturation in YPD to SPO. **B**, RNA-seq, TSS-seq and TES-seq tracks of *IRT2* and *IRT1* in *MAT*a/α 6h in SPO (FW1511) and *MAT*a 4h in SPO (FW1509) cells. Displayed are the signals of normalized reads for both the Watson and Crick strands. Cells were grown as described in a. **C**, *IRT1* and *SCR1* expression as detected by northern blot in control (FW1509) and *irt1-T* strains (FW155). The *irt1-T* strain harbors a transcriptional terminator ~ 200 bp after the *IRT1* TSS. Cells were grown as described in a. **D**, Similar data as in B, except that *IRT2* is highlighted, which demonstrates that *IRT2* is transcribed in *MAT*a cells (4h in SPO). **E,** *IRT1*, *IRT2*, and *IME1* expression in WT *MAT*a cells (FW1509, lanes 1-7), and Δ*irt2*(-188) (FW1210, lanes 8-14) and Δ*irt2*(-246) (FW128, lanes 15-21) mutants as presented in Figure 1e. A blot with adjust signal adjusted signal highlighting *IRT2* expression from the northern blot is also included.

**Figure S2. The act of *IRT2* transcription is required for *IRT1* expression**

**A,** Scheme of the *IME1* locus harboring a transcriptional terminator *(irt2-T)* or a control insert (*irt2-I*) in *IRT2*. **B,** *IRT1*, *IRT2*, and *IME1* expression in WT *MAT*a cells

(FW1509, lanes 1-5), Δ*irt2-T* (FW3596, lanes 6-10) and Δ*irt2-I* (FW4175, lanes 11-15) presented in A, as detected by northern blot. *SCR1* was used as a loading control. The asterisk depicts the prematurely terminated form of *IRT2*. Cells were grown in rich medium (YDP) to saturation, shifted to pre-SPO, and subsequently transferred to SPO. Samples were taken at the indicated time points. **C**, Rme1 chromatin immunoprecipitation followed by quantitative PCR using oligos nested in *IRT2* in no tag control, *RME1-V5* control, *Δirt2(-246)* and *irt1-T* single mutants and *Δirt2(-246) irt1-T* double mutant (FW1509, FW531, FW3140, FW155 and FW8909). Cells were grown as in A and samples were formaldehyde fixed at the indicated time points. Chromatin immunoprecipitation was performed and recovered DNA fragments were quantified by quantitative PCR using a primer pair directed to Rme1 binding sites in the *IRT1* promoter. Signals were normalized to *HMR*, which does not bind Rme1. The error bars represent the standard error of the mean (SEM) of n = 3 biological repeats. *** correspond to a p-value < 0.0005 on a two-way ANOVA analysis followed by a Fisher's LSD test comparing the mutant strains with the control strain. **D,** Western blot of Rpb3-AID and Rme1-V5 expression in IAA treated or mock treated cells. *MAT*a cells harbouring *RPB3-AID* and *RME1-V5* (FW6467) were grown to saturation in rich medium (YPD), treated with IAA to deplete Rpb3-AID (Pol II) and subsequently IAA was washed out from the medium, and cells were transferred to SPO. **E,** Rme1 association at *IRT2* locus in condition described in C, using *MAT*a cells harbouring either *RPB3-AID* or *RME1-V5,* or both (FW5735, FW4031 and FW6467). **** correspond to a p-value < 0.0001 on a two-way ANOVA followed by Fisher's LSD test comparing DMSO and IAA treated cells. **F**, Quantification of *IRT1* expression described in Figure 2H. *IRT1* signals were normalized over *SCR1*. n = 2 +/- SEM. *, ** and *** correspond to p-value < 0.05, <

0.005, < 0.0005 respectively on a two-way ANOVA analysis followed by a Fisher's LSD test. **G,** *IRT1* and *IRT2* expression in *MAT*a cells (FW1509, lanes 1-11) and *u6bs*Δ cells harboring the deletion of the Ume6 binding site in absence of a selectable marker *u6bs*Δ (FW1378, lanes 12-22) or with a selectable marker *u6bsΔ::KanMX* (FW2438, lanes 23-33). *SCR1* was used as a loading control. Cells were grown in rich medium (YPD) to saturation and subsequently transferred to SPO. Samples were taken at the indicated time points. **H,** ChIP of Rme1-V5 at *IRT1* under the same condition describe in figure 2H, except for the presence of the *RME1-V5* allele in *MAT*a WT (FW4031) and *u6bs*Δ (FW3144). n = 4 +/- SEM. * and ** correspond to a p-value < 0.05 and < 0.005 respectively on a two-way ANOVA followed by Fisher's LSD test comparing *u6bs*Δ to the *MAT*a control. **I,** *IRT1* and *IRT2* expression in *MAT*a cells (FW1509, lanes 1-3), *u6bsΔ::KanMX* (FW2438, lanes 4-6), *ime1-t* (FW2189, lanes 7-9) and *u6bsΔ::KanMX ime1-t* double mutant (FW2539, lanes 10-12), detected by northern blot. *SCR1* was used as a loading control. Cells were grown in rich medium (YPD) to saturation, transferred to pre-SPO and grown for an addition 18 hours, and subsequently transferred to SPO medium. Samples were taken at the indicated time points. **J,** *IRT1*, *IRT2*, and *IME1* expression in *MAT*a cells (FW1509, lane 1-5), Δ*irt2-T* (FW3596, lanes 6-10), *u6bsΔ::KanMX* (FW2438, lanes 11-15) and *u6bsΔ::KanMX* Δ*irt2-T* double mutant (FW3493, lanes 16-20) detected by northern blot. *SCR1* was used as a loading control. Cells were grown as in d and samples were taken at the indicated time points.

**Figure S3. *IRT2* prevents entry into gametogenesis in cells with a single mating-type**

**A,** Representative images of *IME1* and *ACT1* expression in single cells as measured by single-molecule RNA FISH in control *MAT*a (FW1534), Δ*irt2*(-246) (FW3580) and *irt2-T* (FW3585). Cells were grown overnight to saturation in rich medium (Y), shifted to pre-SPO and grown for an addition 18 hours, and subsequently transferred to SPO. Cells were fixed at the indicated time points and hybridized with probes directed against *IME1* and *ACT1*. **B,** Distribution of *IME1* expression in single cells. The data described in Figure 3A were binned according to the number of *IME1* copies per cell.

**Figure S4. Rtt109 and Asf1 promote *IRT1* expression**

**A,** *IRT1*, *IRT2*, and *IME1* expression in *MAT*a control (FW1509), *rtt109Δ* (FW4077) and *asf1Δ* (FW4521) cells detected by northern blot. *SCR1* was used as loading control. Cells were grown in rich medium to saturation, shifted and grown in pre-SPO, and subsequently transferred to SPO. Samples were taken at the indicated time points. **B,** Rme1-V5 control *MAT*a cells (FW4031) and *rtt109Δ* mutant (FW4075), as detected by western blot using anti-V5 (green channel). Hxk1 was used as loading controls using anti-Hxk1 (red channel) antibodies. Cells were grown as described in a and samples were taken at the indicated time points. **C**, Nascent RNA-seq for *RPL18B*. Two replicates of RNA-seq and nascent RNA-seq (Pol II associated RNA) are displayed. Cells were harvested in SPO 4h and during exponential growth (log phase YPD) for *MAT*a control, and SPO 4h for *MAT*a *rtt109Δ*. Subsequently, Pol II was purified using the Rpb3-FLAG. The values on the y-axes indicate Reads Per Million (RPM). **D**, Similar as C, except that *IRT1* and *IME1* transcription is displayed. **E**, Quantification of *IRT1* and *IME1* expression levels described in Figure 4E. *IRT1* signals were normalized over *SCR1*. n = 3 +/- SEM. * and ** correspond to p-value < 0.05 and < 0.005 on a two-way ANOVA followed by a Fisher's LSD test. **F,** *IRT1* and *IRT2* expression in control *MAT*a cells (FW1509, lanes 1-5), *u6bs*Δ (FW2438, lanes 6-10), *rtt109*Δ (FW4077, lanes 11-15) and *u6bs*Δ *rtt109*Δ double mutant (FW5225, lanes 16-20). Cells were grown as in A, and samples were taken at the indicated time points. An adjusted signal highlighting *IRT2* expression from the northern blot is also included. **G,** Quantification of *IRT1* expression for data described in F. *IRT1* expression was normalized over *SCR1*. n = 3 +/- SEM. *, ** and **** correspond to p-value < 0.05, < 0.005 and < 0.0001 respectively on a two-way ANOVA followed by a Fisher's LSD test.

**Figure S5. *IRT2* links H3K56ac to nucleosome dynamics**

**A,** Quantification of *IRT1* and *IME1* expression of data shown in Figure 5A. *IRT1* and *IME1* signals were normalized over *SCR1*. n = 3, +/- SEM. *, ** and *** correspond to p-value < 0.05, < 0.005 and < 0.0005 respectively on a two-way ANOVA followed by a Fisher's LSD test. **B**, *IRT1* and *IRT2* expression in control *MAT*a H3 (FW5102, lanes 1-4), H3 *rtt109*Δ (FW6443, lanes 5-8) and *H3K56R* (FW5116, lanes 9-12) and *rtt109*Δ *H3K56R* double mutant (FW5724, lanes 13-16) cells as detected by northern blot. *SCR1* was used as loading control. Same blot with adjusted signal highlighting *IRT2* expression from the northern blot is also included. Cells were grown as in b. Samples were taken at the indicated time points. **C**, Quantification of *IRT1* expression as described in c. *IRT1* signals were normalized to *SCR1*. n = 2, mean values and data range are displayed. **, *** and **** correspond to p-value < 0.005, < 0.0005 and < 0.0001 respectively, on a two-way ANOVA followed by a Fisher's LSD test. **D,** Histone exchange at the *IRT2, PHO5* and *PGK1* promoters. Strains as described in Figure 5C an 5D. To have constitutive levels of *IRT2* transcription we used the *u6bs*Δ mutation (FW7880), while the wild-type cells (FW7853), display no *IRT2* transcription in rich medium (YP). Cells were grown as described in Figure 5D. Samples were taken at the indicated time point for ChIP. Primer pairs for the *PHO5* and *PGK1* promoter were used for the analysis. The signal for each histone H3 ChIP (Myc-H3 and FLAG-H3) were normalized to a telomere locus, and ratio for n = 3 +/- SEM is displayed. The start (0 min) and end points (60 min) of galactose induced and non-induced cells are displayed.

**Figure S6. Distinct levels of *IRT2* lead to opposite outcomes of *IRT1* transcription**

**A**, *IRT1* and *IRT2* expression in control *MAT*a (FW6560, lanes 1-6), $lexO_1$(+96) (FW6594, lanes 7-12), $lexO_2$(+96) (FW6599, lanes 13-18), $lexO_3$(+96) (FW6607, lanes 19-24), $lexO_4$(+96) (FW6611, lanes 25-30), and $lexO_8$(+96) (FW6619, lanes 31-36) cells as detected by northern blot. These cells also expressed LexA-ER. Cells were grown in rich medium (YPD) to saturation, shifted and grown in pre-SPO for 15h, and cells either mock treated or treated with β-estradiol (25nM) for additional 1 hour. Subsequently cells were transferred to SPO and the treatments were continued. Samples were taken at the indicated time points. *SCR1* was used as loading control. **B,** Quantification of *IRT1* expression as described in A. Signals were normalized to *SCR1* and presented as a ratio of β-estradiol over mock treatment for the 1h time point. **C,** Chromatin structure at the *IRT1* promoter in the presence of distinct levels of *IRT2* transcription. *MAT*a control cells (FW6560) or cells harbouring 1 or 4 lexO(+96) sites (FW6594 or FW6611) were grown as described in a and the mock treated condition is displayed. β-estradiol treated condition is displayed in figure 6C. Cells were grown in rich nutrient conditions (YPD) till exponential growth were included in the analysis. Cells were fixed with formaldehyde, and chromatin was digested with micrococcal nuclease (MNase) followed by qPCR using scanning primer pairs in *IRT2*. The red arrows indicate the position of the Rme1 binding sites. The signals were normalized over a telomeric region from chromosome VI. n = 3 plus SEM. **D**, Pol II binding at *IRT1* as detected by ChIP. Similar as C, except that cells also contained Rpb3-FLAG, which was used for the immunoprecipitation step. Cells were grown as described in C, and samples for ChIP were taken at 3h in SPO. Signals were normalized over the silent mating type locus *HMR*. The ratio β-estradiol treated over mock treated are presented. n = 4 +/- SEM. ** and *** correspond to p-value < 0.005 and < 0.0005 respectively on a one-way ANOVA followed by a Fisher's LSD test.

**Figure S7. Modelling of mating-type control of Ime1 expression involving two lncRNAs**

**A,** Model for Ime1 regulation by *IRT1*, *IRT2*, and *IME1* (see Methods section for details). **B,** Simulation of *IRT1*, *IRT2*, and *IME1* transcription rates, *IME1* mRNA and Ime1 protein levels in the presence of different concentrations of Rme1. Indicated are different Rme1 concentrations (r= 0, 0.05, 0.1, 0.25, 0.5 and 5, arbitrary units) leading to either no entry into meiosis in *MAT*a or *MAT*α (haploid), or entry in meiosis in *MAT*a/α (diploid) cells. **C,** Dose response relationship between Ime1 steady state levels and Rme1 concentration simulated for WT *IRT2* (activating and repressing levels) and *IRT2* activating levels only, which are simulated by setting I2=A. Indicated are the levels of Rme1 corresponding to either no entry into meiosis in haploid cells, or entry in meiosis into diploid cells.

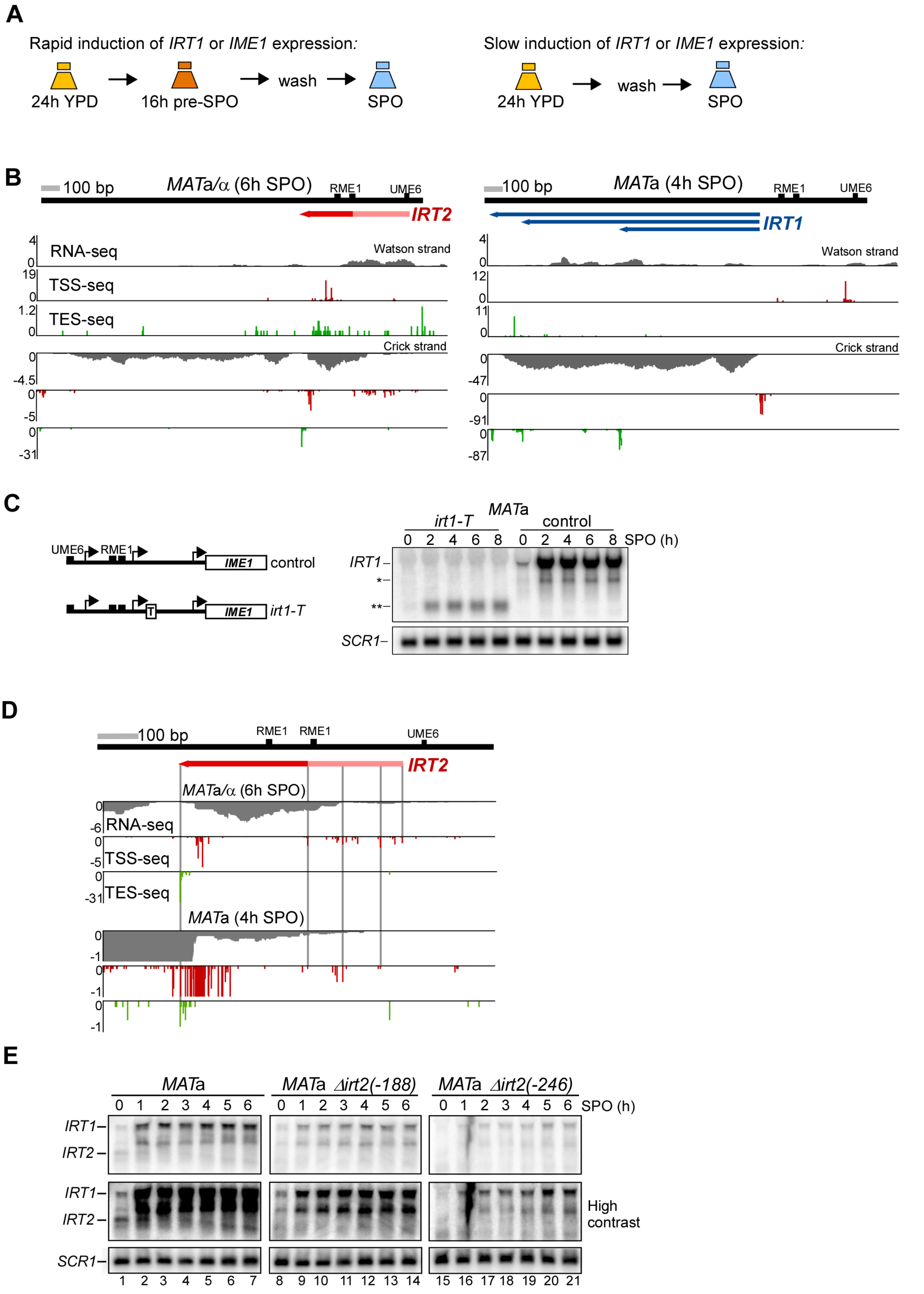
A
Rapid induction of IRT1 or IME1 expression:
24h YPD
16h pre-SPO
wash
SPO
Slow induction of IRT1 or IME1 expression:
24h YPD
wash
SPO
B
100 bp
MATa/α (6h SPO)
RME1
UME6
IRT2
RNA-seq
TSS-seq
TES-seq
Watson strand
Crick strand
100 bp
MATa (4h SPO)
IRT1
C
UME6
RME1
IME1
control
irt1-T
MATa
irt1-T
control
0 2 4 6 8 0 2 4 6 8 SPO (h)
IRT1
*
**
SCR1
D
100 bp
RME1
RME1
UME6
IRT2
MATa/α (6h SPO)
RNA-seq
TSS-seq
TES-seq
MATa (4h SPO)
E
MATa
MATa Δirt2(-188)
MATa Δirt2(-246)
0 1 2 3 4 5 6 SPO (h)
IRT1
IRT2
High contrast
SCR1
1 2 3 4 5 6 7 8 9 10 11 12 13 14 15 16 17 18 19 20 21

Moretto *et al*. Figure S2

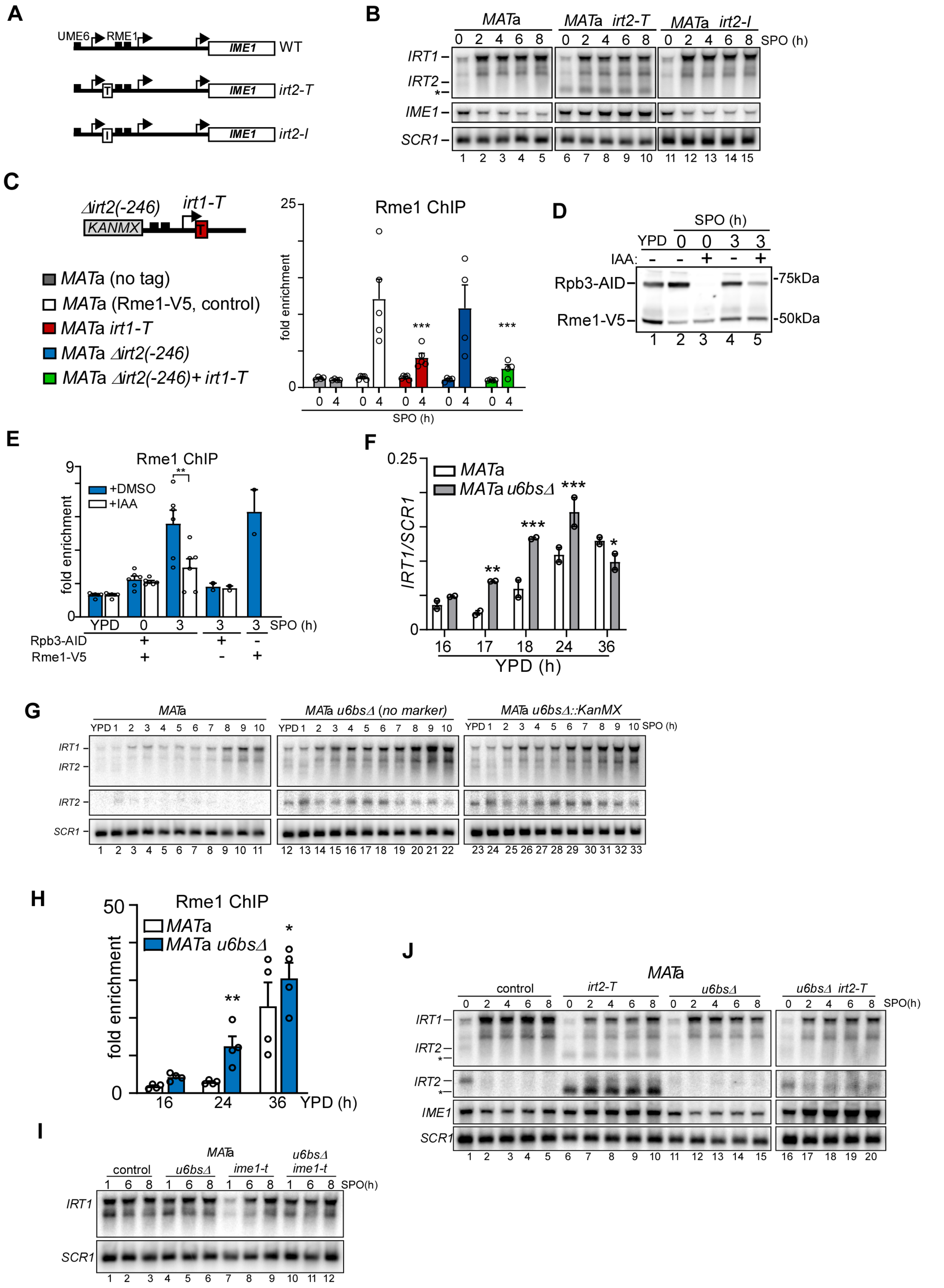

Moretto *et al.* Figure S3

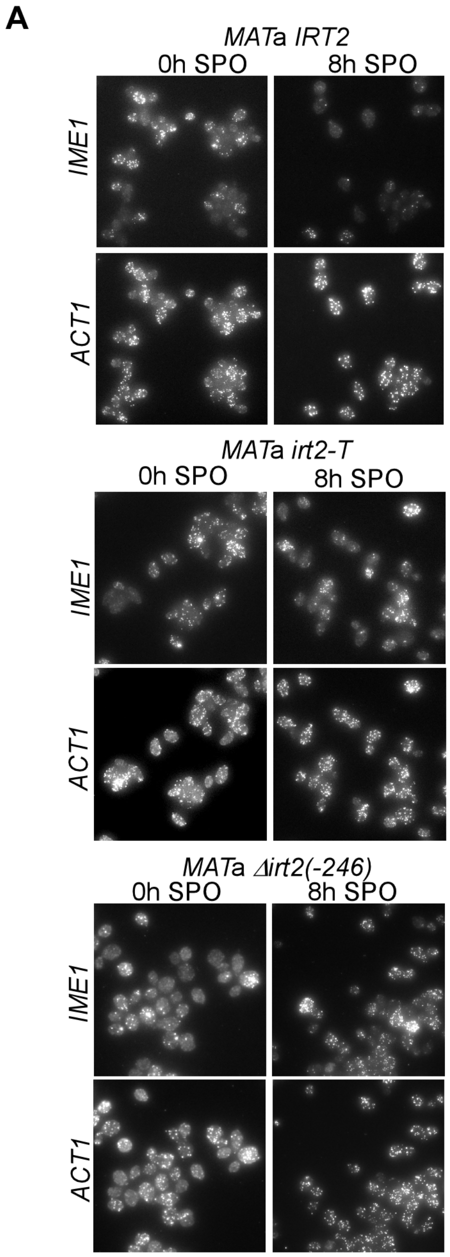


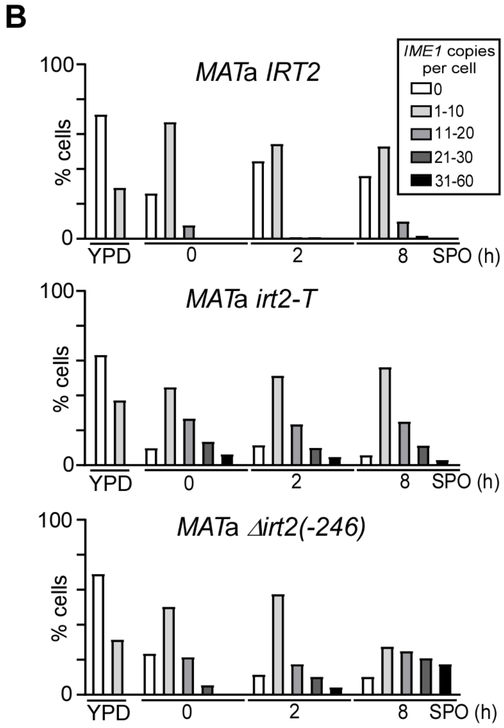

Moretto *et al*. Figure S4

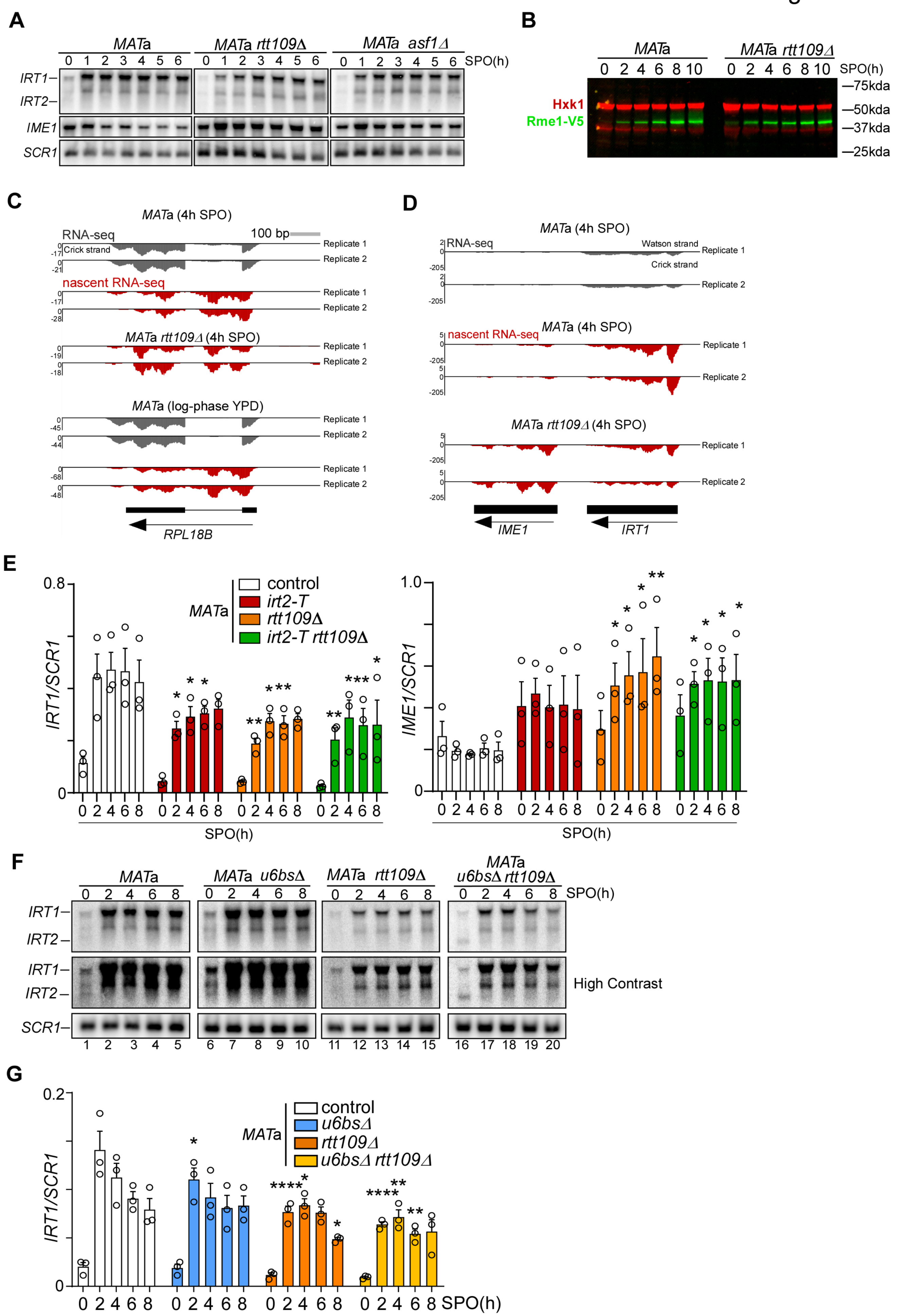

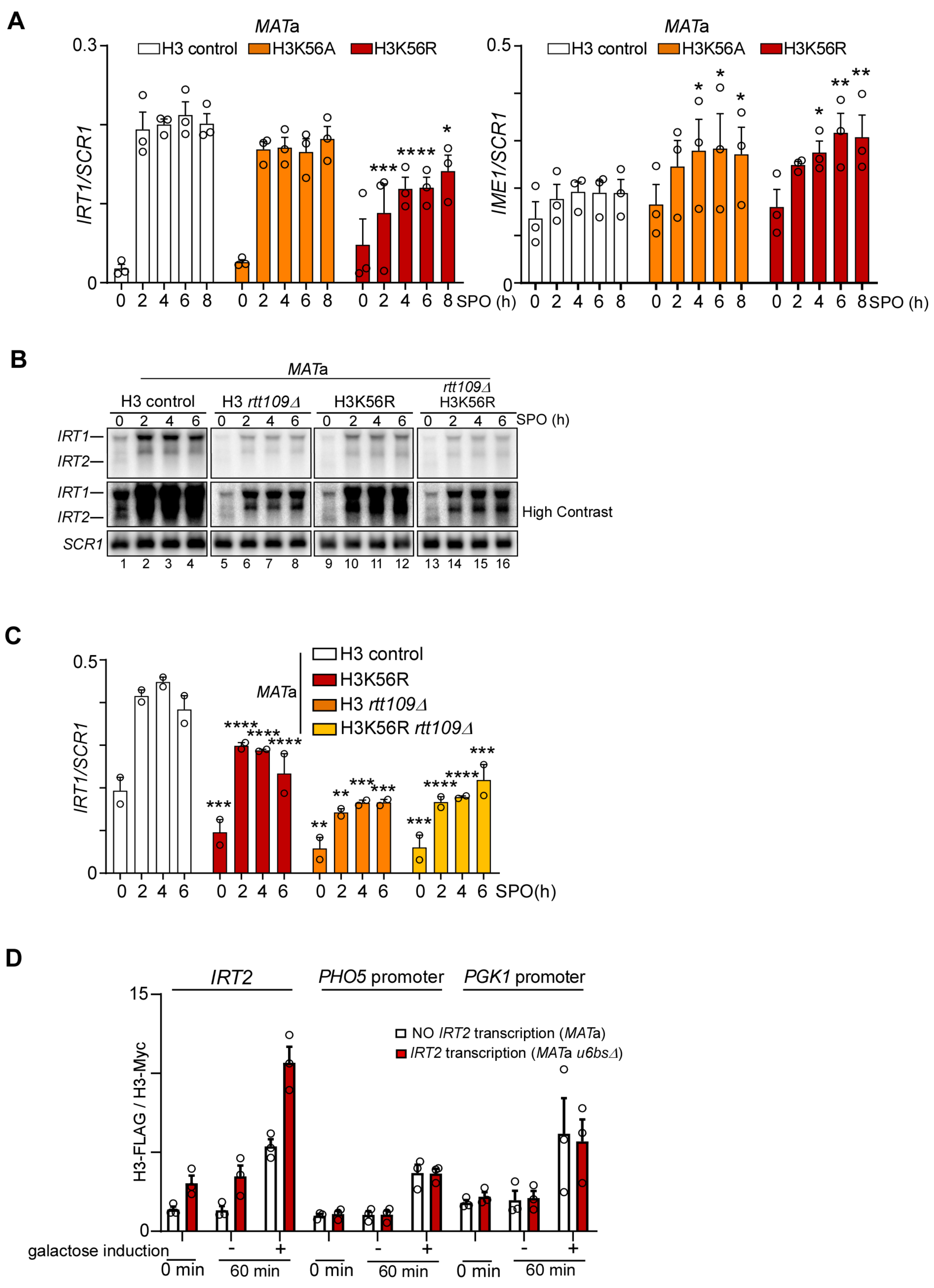
A
MATa
H3 control
H3K56A
H3K56R
IRT1/SCR1
IME1/SCR1
SPO (h)
B
MATa
H3 control
H3 rtt109Δ
H3K56R
rtt109Δ H3K56R
SPO (h)
IRT1
IRT2
High Contrast
SCR1
C
MATa
H3 control
H3K56R
H3 rtt109Δ
H3K56R rtt109Δ
IRT1/SCR1
SPO(h)
D
IRT2
PHO5 promoter
PGK1 promoter
NO IRT2 transcription (MATa)
IRT2 transcription (MATa u6bsΔ)
H3-FLAG / H3-Myc
galactose induction
0 min
60 min

### Moretto *et al.* Figure S6

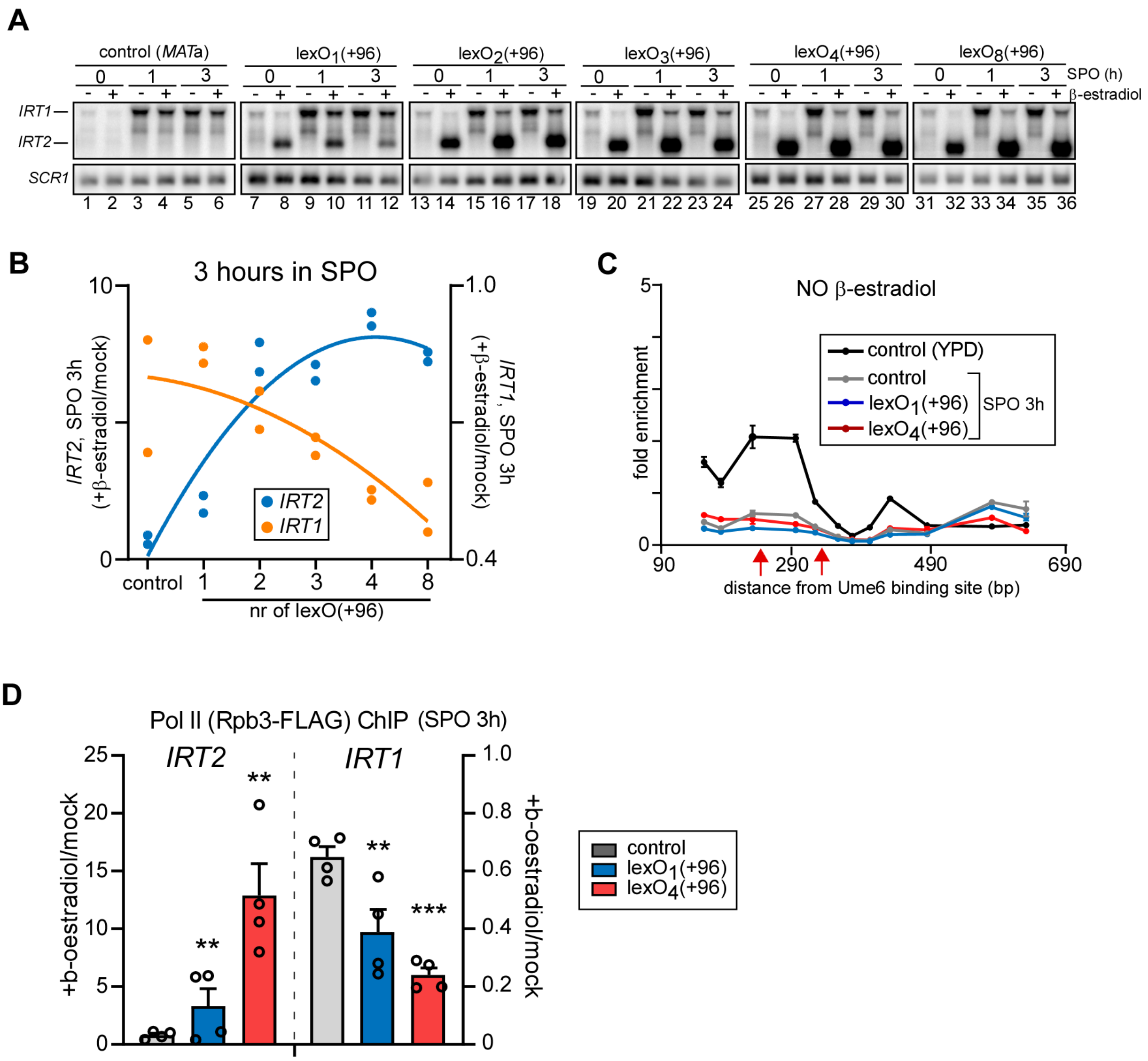

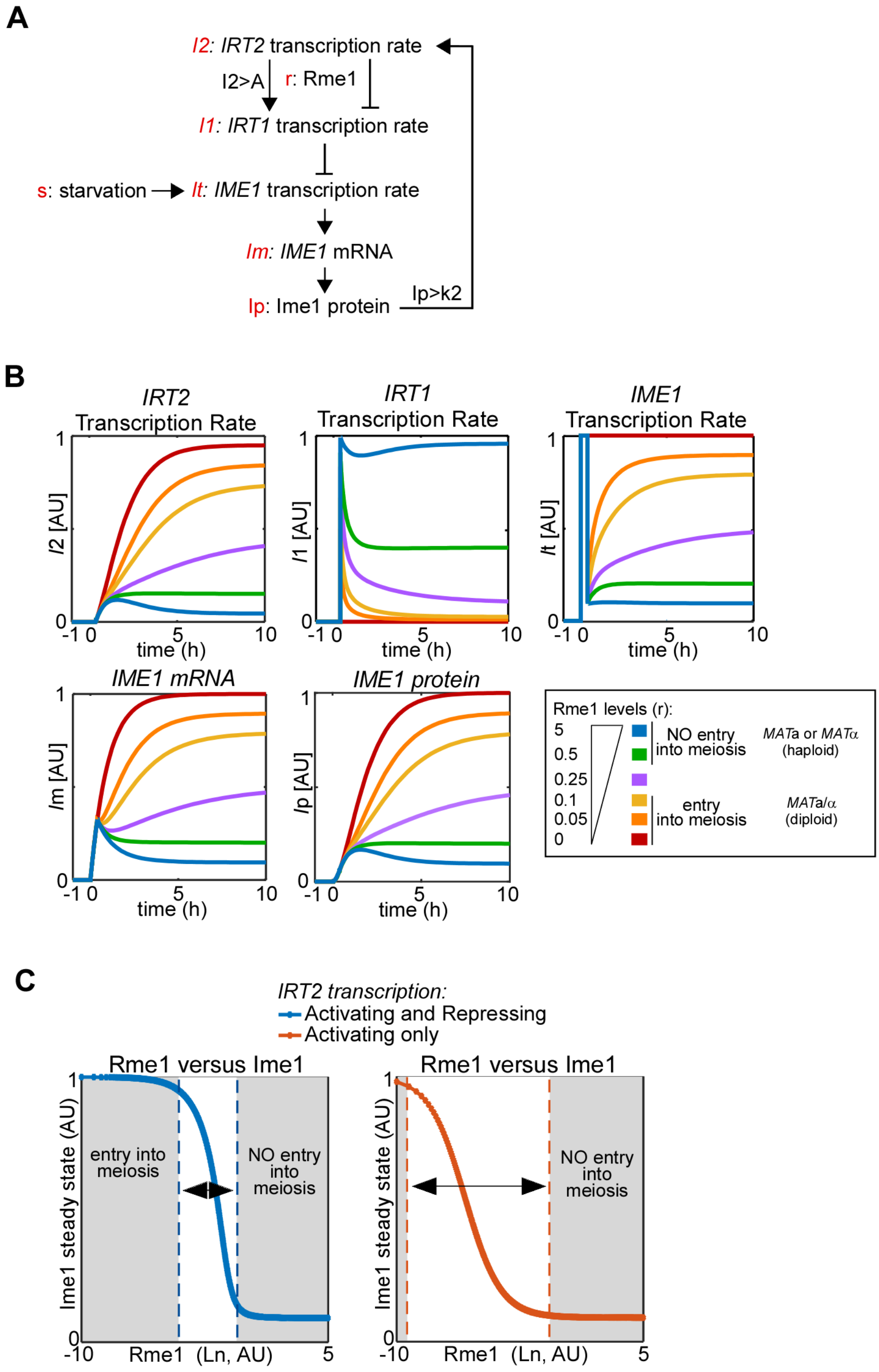
A
I2: IRT2 transcription rate
I2>A
r: Rme1
I1: IRT1 transcription rate
s: starvation
It: IME1 transcription rate
Im: IME1 mRNA
Ip: Ime1 protein
Ip>k2
B
IRT2 Transcription Rate
I2 [AU]
IRT1 Transcription Rate
I1 [AU]
IME1 Transcription Rate
It [AU]
IME1 mRNA
Im [AU]
IME1 protein
Ip [AU]
time (h)
Rme1 levels (r):
5
0.5
0.25
0.1
0.05
0
NO entry into meiosis
MATa or MATα (haploid)
entry into meiosis
MATa/α (diploid)
C
IRT2 transcription:
Activating and Repressing
Activating only
Rme1 versus Ime1
Ime1 steady state (AU)
entry into meiosis
NO entry into meiosis
Rme1 (Ln, AU)

**Table S1. Mutants that affect *IRT1* expression**

| gene delete | *IRT1* | *IME1* |
|---|---|---|
| *ASF1* | - | + |
| *CAC2* | 0 | NT |
| *CTK1* | --- | + |
| *ELP4* | 0 | 0 |
| *ELP6* | 0 | 0 |
| *GCN5* | --- | +++ |
| *HIR1* | 0 | NT |
| *HST3* | 0 | NT |
| *HST4* | 0 | NT |
| *JHD2* | 0 | 0 |
| *NAP1* | 0 | 0 |
| *RLF2* | + | NT |
| *RPB9* | -- | ++ |
| *RTR1* | 0 | 0 |
| *RTT106* | 0 | NT |
| *RTT109* | - | ++ |
| *SET1* | 0 | 0 |
| *SET3* | 0 | + |
| *SNF5* | --- | NT |
| *SUB1* | 0 | 0 |
| *SWC3* | 0 | 0 |
| *SWC5* | 0 | 0 |
| *SWR1* | 0 | NT |
| *VPS75* | 0 | NT |
| *HOS2* | 0 | NT |

**NT: not tested**
**+/-: difference with WT**
**0: no difference with WT**

**Table S2. Genotypes of yeast strains**

| | |
|---|---|
| FW1509 | *MAT*a, *ho::LYS2, lys2, ura3, leu2::hisG, his3::hisG, trp1::hisG* |
| FW1511 | *MAT*a, *ho::LYS2, lys2, ura3, leu2::hisG, his3::hisG, trp1::hisG*<br>*MAT*alpha, *ho::LYS2, lys2, ura3, leu2::hisG, his3::hisG, trp1::hisG* |
| FW1356 | *MAT*a, *ho::LYS2, ura3, leu2::hisG, his3::hisG, trp1::hisG,*<br>*irt2Δ(-246bp)::KanMX* |
| FW1210 | *MAT*a, *ho::LYS2, ura3, leu2::hisG, his3::hisG, trp1::hisG,*<br>*irt2Δ(-188bp)::HisMX* |
| FW3596 | *MAT*a, *ho::LYS2, lys2, ura3, leu2::hisG, his3::hisG, trp1::hisG*<br>*irt2-Term* |
| FW4175 | *MAT*a, *ho::LYS2, lys2, ura3, leu2::hisG, his3::hisG, trp1::hisG*<br>*IRT2-Insert* |
| FW4031 | *MAT*a, *ho::LYS2, lys2, ura3, leu2::hisG, his3::hisG, trp1::hisG,*<br>*RME1::3xV5::HisMX* |
| FW3132 | *MAT*a, *ho::LYS2, lys2, ura3, leu2::hisG, his3::hisG, trp1::hisG,*<br>*RME1::3xV5::HisMX*<br>*irt2Δ(-188bp)::HisMX* |
| FW3140 | *MAT*a, *ho::LYS2, lys2, ura3, leu2::hisG, his3::hisG, trp1::hisG,*<br>*RME1::3xV5::HisMX*<br>*irt2Δ(-246bp)::KanMX* |
| FW3128 | *MAT*a, *ho::LYS2, lys2, ura3, leu2::hisG, his3::hisG, trp1::hisG*<br>*irt2-Term*<br>*RME1::3xV5::HisMX* |
| FW6467 | *MAT*a, *ho::LYS2, lys2, ura3, leu2::hisG, his3::hisG, trp1::hisG*<br>*his3:: pCUP-OsTIR-HIS3*<br>*RBP3:3V5-IAA17:KanMX6*<br>*RME1::3xV5::HisMX* |
| FW5735 | *MAT*a, *ho::LYS2, lys2, ura3, leu2::hisG, his3::hisG, trp1::hisG*<br>*his3:: pCUP-OsTIR-HIS3*<br>*RBP3::3V5-IAA17::KanMX6* |
| FW2189 | *MAT*a, *ho::LYS2, lys2, ura3, leu2::hisG, his3::hisG, trp1::hisG*<br>*IME1-NatMX4* |
| FW2438 | *MAT*a, *ho::LYS2, lys2, ura3, leu2::hisG, his3::hisG, trp1::hisG*<br>*u6bsΔ::KanMX* |
| FW2539 | *MAT*a, *ho::LYS2, lys2, ura3, leu2::hisG, his3::hisG, trp1::hisG*<br>*Ime1-NatMX4*<br>*u6bsΔ::KanMX* |
| FW3493 | *MAT*a, *ho::LYS2, lys2, ura3, leu2::hisG, his3::hisG, trp1::hisG*<br>*u6bsΔ::KanMX*<br>*irt2-Term* |
| FW1378 | *MAT*a, *ho::LYS2, lys2, ura3, leu2::hisG, his3::hisG, trp1,*<br>*u6bsΔ* |
| FW3144 | *MAT*a, *ho::LYS2, lys2, ura3, leu2::hisG, his3::hisG, trp1,*<br>*RME1::3xV5::HisMX*<br>*u6bsΔ::KanMX* |
| FW3585 | *MAT*a, *ho::LYS2, lys2, ura3, leu2::hisG, his3::hisG, trp1::hisG*<br>*flo8::KanMX6*<br>*irt2-Term* |

| | |
|---|---|
| FW3580 | *MAT*a*, ho::LYS2, lys2, ura3, leu2::hisG, his3::hisG, trp1::hisG*<br>*flo8::KanMX6*<br>*irt2Δ(-246bp)::KanMX* |
| FW1533 | *MAT*a*, ho::LYS2, lys2, ura3, leu2::hisG, his3::hisG, trp1::hisG*<br>*flo8::KanMX6* |
| FW15 | *MAT*a*, ho::LYS2, lys2, ura3, leu2::hisG, his3::hisG, trp1::hisG*<br>*matalpha::MAT*a*::URA3, ho::LYS2, lys2, ura3, leu2::hisG, his3::hisG, trp1::hisG* |
| FW2342 | *MAT*alpha*, ho::LYS2, lys2, ura3, leu2::hisG, his3::hisG, trp1::hisG,*<br>*rme1::HIS3*<br>*MAT*a*, ho::LYS2, lys2, ura3, leu2::hisG, his3::hisG, trp1::hisG,*<br>*rme1::HIS3* |
| FW3402 | *MAT*a*, ho::LYS2, lys2, ura3, leu2::hisG, his3::hisG, trp1::hisG*<br>*irt2Δ(-246bp)::KanMX*<br>*matalpha::MAT*a*::URA3, ho::LYS2, lys2, ura3, leu2::hisG, his3::hisG, trp1::hisG*<br>*irt2Δ(-246bp)::KanMX* |
| FW3453 | *MAT*a*, ho::LYS2, ura3, leu2::hisG, his3::hisG, trp1::hisG,*<br>*irt2Δ(-188bp)::HisMX*<br>*matalpha::MAT*a*::URA3, ho::LYS2, ura3, leu2::hisG, his3::hisG, trp1::hisG,*<br>*irt2Δ(-188bp)::HisMX* |
| FW1317 | *MAT*a*, ho::LYS2, lys2, ura3, leu2::hisG, his3::hisG, trp1::hisG,*<br>*rme1Δ::HisMx*<br>*matalpha::MAT*a*::URA3, ho::LYS2, lys2, ura3, leu2::hisG, his3::hisG, trp1::hisG,*<br>*rme1Δ::HisMx* |
| FW3629 | *MAT*a*, ho::LYS2, lys2, ura3, leu2::hisG, his3::hisG, trp1::hisG,*<br>*irt2-Term*<br>*matalpha::MAT*a*::URA3, ho::LYS2, lys2, ura3, leu2::hisG, his3::hisG, trp1::hisG,*<br>*irt2-Term* |
| FW4077 | *MAT*a*, ho::LYS2, lys2, ura3, leu2::hisG, his3::hisG, trp1::hisG*<br>*rtt109:HygMx* |
| FW5225 | *MAT*a*, ho::LYS2, lys2, ura3, leu2::hisG, his3::hisG, trp1::hisG*<br>*rtt109:HygMx*<br>*u6bsΔ::KanMX* |
| FW4072 | *MAT*alpha*, ho::LYS2, lys2, ura3, leu2::hisG, his3::hisG, trp1::hisG*<br>*rtt109:HygMx*<br>*irt2-Term* |
| FW4075 | *MAT*a*, ho::LYS2, lys2, ura3, leu2::hisG, his3::hisG, trp1::hisG*<br>*rtt109:HygMx*<br>*RME1::3xV5::HisMX* |
| FW4073 | *MAT*a*, ho::LYS2, lys2, ura3, leu2::hisG, his3::hisG, trp1::hisG*<br>*rtt109:HygMx*<br>*RME1::3xV5::HisMX*<br>*irt2-Term* |
| FW5102 | *MAT*a*, ho::LYS2, lys2, ura3, leu2::hisG, trp1::hisG, his4-N/G*<br>*hhf1-hht1::LEU2 hhf2-hht2::trp1::KanMX*<br>*plasmid FW502 (HHT1-HHF1 CEN ARS URA3)* |
| FW5116 | *MAT*a*, ho::LYS2, lys2, ura3, leu2::hisG, trp1::hisG, his4-N/G*<br>*hhf1-hht1::LEU2 hhf2-hht2::trp1::KanMX*<br>*plasmid FW504 (HHT1-HHF1 H3k56R CEN ARS URA3)* |
| FW5113 | *MAT*a*, ho::LYS2, lys2, ura3, leu2::hisG, trp1::hisG, his4-N/G*<br>*hhf1-hht1::LEU2 hhf2-hht2::trp1::KanMX*<br>*plasmid FW503 (HHT1-HHF1 H3k56A CEN ARS URA3)* |

| | |
|---|---|
| FW7413 | MATa, ho::LYS2, lys2, ura3, leu2::hisG, trp1::hisG, his4-N/G<br>hhf1-hht1::LEU2 hhf2-hht2::trp1::KanMX<br>matalpha::TRP1-MATa, ho::LYS2, lys2, ura3, leu2::hisG, trp1::hisG, his4-N/G<br>hhf1-hht1::LEU2 hhf2-hht2::trp1::KanMX<br>plasmid FW502 (HHT1-HHF1 CEN ARS URA3) |
| FW7417 | MATa, ho::LYS2, lys2, ura3, leu2::hisG, trp1::hisG, his4-N/G<br>hhf1-hht1::LEU2 hhf2-hht2::trp1::KanMX<br>matalpha::TRP1-MATa, ho::LYS2, lys2, ura3, leu2::hisG, trp1::hisG, his4-N/G<br>hhf1-hht1::LEU2 hhf2-hht2::trp1::KanMX<br>plasmid FW504 (HHT1-HHF1 H3k56R CEN ARS URA3) |
| FW7508 | MATa, ho::LYS2, lys2, ura3, leu2::hisG, trp1::hisG, his4-N/G<br>hhf1-hht1::LEU2 hhf2-hht2::trp1::KanMX<br>matalpha::TRP1-MATa, ho::LYS2, lys2, ura3, leu2::hisG, trp1::hisG, his4-N/G<br>hhf1-hht1::LEU2 hhf2-hht2::trp1::KanMX<br>plasmid FW503 (HHT1-HHF1 H3k56A CEN ARS URA3) |
| FW6443 | MATa, ho::LYS2, lys2, ura3, leu2::hisG, trp1::hisG, his4-N/G<br>hhf1-hht1::LEU2 hhf2-hht2::trp1::KanMX<br>plasmid FW502 (HHT1-HHF1 CEN ARS URA3)<br>rtt109:HygMx |
| FW5724 | MATa, ho::LYS2, lys2, ura3, leu2::hisG, trp1::hisG, his4-N/G<br>hhf1-hht1::LEU2 hhf2-hht2::trp1::KanMX<br>plasmid FW504 (HHT1-HHF1 H3k56R CEN ARS URA3)<br>rtt109:HygMx |
| FW1555 | MATa, ho::LYS2, lys2, ura3, leu2::hisG, his3::hisG, trp1::hisG,<br>ime1::HisMX6 |
| FW7142 | MATa, ho::LYS2, lys2, ura3, leu2::hisG, his3::hisG, trp1::hisG,<br>ime1::HISMx6<br>KanMX::$LexO_1$(-10)::IRT2 |
| FW6560 | MATa, ho::LYS2, lys2, ura3, leu2::hisG, his3::hisG, trp1::hisG,<br>pGDP1::LexA-ER-HA-B112::TRP1 |
| FW6594 | MATa, ho::LYS2, lys2, ura3, leu2::hisG, his3::hisG, trp1::hisG,<br>pGDP1::LexA-ER-HA-B112::TRP1<br>KanMX::$LexO_1$(+96)::IRT2 |
| FW6599 | MATa, ho::LYS2, lys2, ura3, leu2::hisG, his3::hisG, trp1::hisG,<br>pGDP1::LexA-ER-HA-B112::TRP1<br>KanMX::$LexO_2$(+96)::IRT2 |
| FW6607 | MATa, ho::LYS2, lys2, ura3, leu2::hisG, his3::hisG, trp1::hisG,<br>pGDP1::LexA-ER-HA-B112::TRP1<br>KanMX::$LexO_3$(+96)::IRT2 |
| FW6611 | MATa, ho::LYS2, lys2, ura3, leu2::hisG, his3::hisG, trp1::hisG,<br>pGDP1::LexA-ER-HA-B112::TRP1<br>KanMX:: $LexO_4$(+96)::IRT2 |
| FW6619 | MATa, ho::LYS2, lys2, ura3, leu2::hisG, his3::hisG, trp1::hisG,<br>pGDP1::LexA-ER-HA-B112::TRP1<br>KanMX::$LexO_8$(+96)::IRT2 |
| FW4521 | MATa, ho::LYS2, lys2, ura3, leu2::hisG, his3::hisG, trp1::hisG<br>asf1::HygMx |
| FW7853 | MATa, ura3–52, lys2–801, ade2–101, trp1Δ63, his3Δ200, leu2Δ1<br>hht1-hhf1::pWZ405-F2F9-LEU2<br>hht2-hhf2::pWZ403-F4F10-HIS3<br>ura3::YIplac211[pGAL1/10 HHF1 FLAG-HHT1]<br>CEN plasmid pNOY439[pHHT2::MYC-HHT2 pHHF2::HHF2] |

| | |
|---|---|
| FW7880 | *MATa, ura3–52, lys2–801, ade2–101, trp1Δ63, his3Δ200, leu2Δ1*<br>*u6bs::KanMX*<br>*hht1-hhf1::pWZ405-F2F9-LEU2*<br>*hht2-hhf2::pWZ403-F4F10-HIS3*<br>*ura3::YIplac211[pGAL1/10 HHF1 FLAG-HHT1]*<br>*CEN plasmid pNOY439[pHHT2::MYC-HHT2 pHHF2::HHF2]* |
| FW155 | *MATa, ho::LYS2, ura3, leu2::hisG, his3::hisG, trp1::hisG, irt1-T* |
| FW4557 | *MATa, ho::LYS2, lys2, ura3, leu2::hisG, his3::hisG, trp1::hisG*<br>*rtt109:HygMx*<br>*MATa::URA, ho::LYS2, lys2, ura3, leu2::hisG, his3::hisG, trp1::hisG*<br>*rtt109:HygMx* |
| FW532 | *MATa, ho::LYS2, ura3, leu2::hisG, his3::hisG, trp1::hisG, RME1::3xV5::HisMX, irt1-T* |
| FW8909 | *MATa, ho::LYS2, ura3, leu2::hisG, his3::hisG, trp1::hisG,*<br>*irt1-T, irt2Δ(-246bp)::KanMX, RME1::3xV5::HisMX* |
| FW8758 | *MATa, ho::LYS2, lys2, ura3, leu2::hisG, his3::hisG, trp1::hisG*<br>*ura3::pGPD1-GAL4(848).ER::URA3, KanMX::pGAL1(-10)::IRT2, ime1::HISMx6* |
| FW8759 | *MATa, ho::LYS2, lys2, ura3, leu2::hisG, his3::hisG, trp1::hisG*<br>*KanMX::pGAL1(-10)::IRT2, ime1::HISMx6* |
| FW8555 | *MATa, ho::LYS2, lys2, ura3, leu2::hisG, his3::hisG, trp1::hisG,*<br>*KanMX::pGAL1(-10)::IRT2, ime1::HISMx6*<br>*rtt109:HygMx* |
| FW8510 | *MATa, ho::LYS2, lys2, ura3, leu2::hisG, his3::hisG, trp1::hisG*<br>*Rpb3-FLAG::NATmx, irt2Δ(-188bp)::HisMX* |
| FW8512 | *MATa, ho::LYS2, lys2, ura3, leu2::hisG, his3::hisG, trp1::hisG*<br>*Rpb3-FLAG::NATmx, irt2-Term* |
| FW8515 | *MATa, ho::LYS2, lys2, ura3, leu2::hisG, his3::hisG, trp1::hisG*<br>*Rpb3-FLAG::NATmx* |
| FW8561 | *MATa, ho::LYS2, lys2, ura3, leu2::hisG, his3::hisG, trp1::hisG*<br>*Rpb3-FLAG::NATmx, rtt109::HygMX* |
| FW6467 | *MATa, ho::LYS2, lys2, ura3, leu2::hisG, his3::hisG, trp1::hisG*<br>*his3:: pCUP-OsTIR-HIS3*<br>*RBP3:3V5-IAA17:KanMX6*<br>*RME1::3xV5::HisMX* |
| FW5735 | *MATa, ho::LYS2, lys2, ura3, leu2::hisG, his3::hisG, trp1::hisG*<br>*his3:: pCUP-OsTIR-HIS3*<br>*RBP3::3V5-IAA17::KanMX6* |

**Table S3: Oligo nucleotide sequences used**

| | | |
|---|---|---|
| FW463 | caacgcctccgataatgtatatg | IME1 northern probe |
| FW464 | acgtcgaaggcaatttctaatg | IME1 northern probe |
| FW481 | atttttagcgactgccgaaa | IRT2 qPCR (Rme1 ChIP) |
| FW482 | atgcaacgcctacttgtttt | IRT2 qPCR (Rme1 ChIP) |
| FW1895 | aaaatgaaaggcagaagatg | IRT2 qPCR (H3 ChIP oligo pair 1) |
| FW1896 | ctggtatggtattgtaagga | IRT2 qPCR (H3 ChIP oligo pair 1) |
| FW1899 | atcatgctgttctttccgcc | IRT2 qPCR (H3 ChIP oligo pair 2) |
| FW1900 | cccacccttcttttattgag | IRT2 qPCR (H3 ChIP oligo pair 2) |
| FW1905 | tcaagaagtccactaaatgg | IRT2 qPCR (H3 ChIP oligo pair 3) |
| FW1906 | acaattttatgcttttgagg | IRT2 qPCR (H3 ChIP oligo pair 3) |
| FW43 | acgatccccgtccaagttatg | HMR qPCR (Rme1 and Pol II ChIP) |
| FW50 | cttcaaaggagtcttaatttccctg | HMR qPCR (Rme1 and Pol II ChIP) |
| FW490 | tgatatgtatgggttaaaaaggatg | IRT1-IRT2 dual northern probe |
| FW540 | ggcagttcaaaggcttttctta | IRT1-IRT2 dual northern probe, ChIP and RT-qPCR, IRT1 |
| FW489 | atgcaacgcctacttgtttt | IRT2 northern probe |
| FW493 | gatggagggttggcataaaa | IRT2 northern probe |
| FW1555 | gctgcagaacttggttcataca | IME1 northern probe |
| FW464 | acgtcgaaggcaatttctaatg | IME1 northern probe |
| FW1841 | gaagtgtcccggctataataaa | SCR1 northern probe |
| FW1842 | gacgctggataaaactcccc | SCR1 northern probe |
| FW2448 | tccgaacgctattccagaaagt | ChIP/MNase normalisation (Telomere locus) |
| FW2449 | ccataatgcctcctatatttagcccttt | ChIP/MNase normalisation (Telomere locus) |
| FW2450 | catatattatctatatcatgc | MNase qPCR |
| FW2451 | ccgtgagggaatacattaat | MNase qPCR |
| FW1899 | atcatgctgttctttccgcc | MNase qPCR |
| FW1900 | cccacccttcttttattgag | MNase qPCR |
| FW1901 | attaatgtattccctcacgg | MNase qPCR |
| FW1902 | cttcttgaggttcttttgacatc | MNase qPCR |
| FW1087 | ggatgtcaaaagaacctcaaga | MNase qPCR |
| FW1088 | tttcggcagtcgctaaaaat | MNase qPCR |
| FW1905 | tcaagaagtccactaaatgg | MNase qPCR |
| FW1906 | acaattttatgcttttgagg | MNase qPCR |
| FW1907 | gccgaaaacgtacggctaac | MNase qPCR |
| FW488 | atttttagcgactgccgaaa | MNase qPCR |
| FW487 | tttttgttatctgcctgaaacg | MNase qPCR |
| FW1908 | gctcactttttttcctaccca | MNase qPCR |

| | | |
|---|---|---|
| FW2452 | gctgtacctcaaaagcataa | MNase qPCR |
| FW2453 | tccagaaacggtttcttatat | MNase qPCR |
| FW1909 | aaaacaagtaggcgttgcat | MNase qPCR |
| FW1910 | accctatttcttcacgaggg | MNase qPCR |
| FW1030 | gagcgccaacactatataag | MNase qPCR |
| FW1031 | caaattctttaaactaagcgc | MNase qPCR |
| FW1913 | gcgcttagtttaaagaatttg | MNase qPCR |
| FW1914 | cctttgttttctctttatcccc | MNase qPCR |
| FW1915 | cttgattattggcattccgc | MNase qPCR |
| FW1916 | ttgctcggaggtactagtca | MNase qPCR |
| FW539 | gggtcttaaatacgcagggaat | ChIP and RT- qPCR, IRT1 |
| FW106 | gtaccaccatgttcccaggtatt | RT- qPCR, ACT1 control |
| FW107 | caagatagaaccaccaatccaga | RT- qPCR, ACT1 control |

**Table S4: Plasmids**

| | | |
|---|---|---|
| pFW7 | pRS306-MATa-URA3 | (van Werven et al., 2012) |
| pFW244 | pRS304 MATa-TRP1 | This study |
| pFW355 | pRS304-GPDpr-CRE-EBD78-CYC1t | Gift Celine Bouchoux |
| pFW211 | pUG6-Myc-C-Avitag | (van Werven and Timmers, 2006) |
| pFW502 | pDM9-HHT2-HHF2-URA3 | (Sommermeyer et al., 2013) |
| pFW503 | pDM9-H3K56A-HFF1-URA3 | This study |
| pFW504 | pDM9-H3K56R-HFF1-URA3 | This study |
| pFW643 | pUB921_p1LexOCYC1-3V5 | Gift Elçin Ünal (Chia et al., 2017) |
| pFW644 | pUB922_p2LexOCYC1-3V5 | Gift Elçin Ünal (Chia et al., 2017) |
| pFW645 | pUB923_p3LexOCYC1-3V5 | Gift Elçin Ünal (Chia et al., 2017) |
| pFW646 | pUB924_p4LexOCYC1-3V5 | Gift Elçin Ünal |
| pFW647 | pUB925_p8LexOCYC1-3V5 | Gift Elçin Ünal (Chia et al., 2017) |
